# Is it Time to Switch Your T1W Sequence? Assessing the Impact of Prospective Motion Correction on the Reliability and Quality of Structural Imaging

**DOI:** 10.1101/666289

**Authors:** Lei Ai, R. Cameron Craddock, Nim Tottenham, Jonathan P Dyke, Ryan Lim, Stanley Colcombe, Michael Milham, Alexandre R. Franco

## Abstract

New large neuroimaging studies, such as the Adolescent Brain Cognitive Development study (ABCD) and Human Connectome Project (HCP) Development studies are adopting a new T1-weighted imaging sequence with prospective motion correction (PMC) in favor of the more traditional 3-Dimensional Magnetization-Prepared Rapid Gradient-Echo Imaging (MPRAGE) sequence. Here, we used a developmental dataset (ages 5-21, N=348) from the Healthy Brain Network (HBN) Initiative to directly compare two widely used MRI structural sequences: one based on the Human Connectome Project (MPRAGE) and another based on the ABCD study (MPRAGE+PMC). We aimed to determine if the morphometric measurements obtained from both protocols are equivalent or if one sequence has a clear advantage over the other. The sequences were also compared through quality control measurements. Inter- and intra-sequence reliability were assessed with another set of participants (N=71) from HBN that performed two MPRAGE and two MPRAGE+PMC sequences within the same imaging session, with one MPRAGE (MPRAGE1) and MPRAGE+PMC (MPRAGE+PMC1) pair at the beginning of the session and another pair (MPRAGE2 and MPRAGE+PMC2) at the end of the session. Intraclass correlation coefficients (ICC) scores for morphometric measurements such as volume and cortical thickness showed that intra-sequence reliability is the highest with the two MPRAGE+PMC sequences and lowest with the two MPRAGE sequences. Regarding inter-sequence reliability, ICC scores were higher for the MPRAGE1 - MPRAGE+PMC1 pair at the beginning of the session than the MPRAGE1 - MPRAGE2 pair, possibly due to the higher motion artifacts in the MPRAGE2 run. Results also indicated that the MPRAGE+PMC sequence is robust, but not impervious, to high head motion. For quality control metrics, the traditional MPRAGE yielded better results than MPRAGE+PMC in 5 of the 8 measurements. In conclusion, morphometric measurements evaluated here showed high inter-sequence reliability between the MPRAGE and MPRAGE+PMC sequences, especially in images with low head motion. We suggest that studies targeting hyperkinetic populations use the MPRAGE+PMC sequence, given its robustness to head motion and higher reliability scores. However, neuroimaging researchers studying non-hyperkinetic participants can choose either MPRAGE or MPRAGE+PMC sequences, but should carefully consider the apparent tradeoff between relatively increased reliability, but reduced quality control metrics when using the MPRAGE+PMC sequence.

## Introduction

New technologies are constantly being developed to improve the quality of Magnetic Resonance Imaging (MRI) sequences. While generally welcomed, such advances can present a significant challenge to longitudinal studies, as well as large-scale data acquisitions, both of which tend to be wary of changing methods mid-study in virtue of the potential introduction of confounds. In light of this, choosing optimal and robust MRI pulse sequences for a study is always a challenging task for a neuroimaging researcher. Since its development in the early 1990s, the T1 weighted 3-Dimensional Magnetization-Prepared Rapid Gradient-Echo Imaging (3D MPRAGE, MPRAGE, or MPR) (Mugler and Brookeman 1990; Brant-Zawadzki, Gillan, and Nitz 1992) has become one of the most widely used MRI sequence by neuroimaging researchers. This sequence, or similar sequences from other manufacturers^1^, has been broadly adopted for studies with large or small sample sizes. However, as with all MRI sequences, it is susceptible to head motion which can significantly alter the quality of the morphometry measurements that are extracted (Reuter et al. 2015; Pardoe, Kucharsky Hiess, and Kuzniecky 2016; A. Alexander-Bloch et al. 2016). In recent years, the MPRAGE sequence has been expanded to include volumetric navigators (vNav), allowing prospective motion correction (PMC) during the acquisition (Tisdall et al. 2012, 2016). These structural sequences with navigator-based PMC have the potential to be transformative for studies involving hyperkinetic populations, such as children, the elderly, or patients with movement disorders. In particular, new large multisite studies are adopting these structural scans with PMC (see Table 1). However, the impact of the change from the traditional MPRAGE sequence to the new MPRAGE sequence with PMC has not been fully quantified (Harms et al. 2018), in part, because few datasets contain a large enough sample size with and without PMC images in the same subjects.

**Table 1.** T1-weighted structural imaging parameters across large imaging studies. Only reporting imaging sequences performed on 3T Siemens MRIs. * We were unable to find all the sequence parameters for the BCP. However, the researchers of BCP state they attempt to match as much as possible the imaging parameters of the other HCP studies (Howell et al. 2019). ** For the HBN study, currently only one of the imaging sites (Citigroup Biomedical Imaging Center CBIC) is collecting two structural scans for all subjects.

| Study | Pulse sequence | Voxel size (mm) | Matrix Size | Num Slices | TI (ms) | TR (ms) | Bandwidth (Hz/Pz) | Parallel Imaging | Partial Fourier | Flip Angle (degrees) | Scanner |
| --- | --- | --- | --- | --- | --- | --- | --- | --- | --- | --- | --- |
| Rockland Sample | MPRAGE | 1.0x1.0x1.0 | 256x256 | 176 | 900 | 1900 | 170 | 2 | off | 9 | Tim-Trio |
| UK Biobank | MPRAGE | 1.0x1.0x1.0 | 256x256 | 208 | 880 | 2000 | 240 | 2 | off | 8 | Siemens 3T Skyra |
| ADNI3 | MPRAGE | 1.0x1.0x1.0 | 256x240 | 176 | 900 | 2300 | 240 | 2 | off | 9 | Many |
| WU-Minn HCP [a.k.a HCP-YA] | MPRAGE | 0.7x0.7x0.7 | 320x320 | 256 | 1000 | 2400 | 210 | 2 | off | 8 | Custom HCP Skyra |
| HCP Aging (HCP-A)<br>And Development (HCP-D) | MPRAGE | 0.8x0.8x0.8 | 320x300 | 208 | 1000 | 2500 | 220 | 2 | off | 8 | Prisma |
|  | MPRAGE +PMC | 0.8x0.8x0.8 | 320x300 | 208 | 1000 | 2500 | 740 | 2 | 6/8 (slice partial Fourier) off for phase | 8 | Prisma |
| HCP Baby (BCP)* | MPRAGE | 0.8x0.8x0.8 | 320x320 | 208 | 1060 | 2400 | - | - | - | 8 | Prisma |
| ABCD | MPRAGE+PMC | 1.0x1.0x1.0 | 256x256 | 176 | 1060 | 2500 | 240 | 2 | off | 8 | Prisma |
| Healthy Brain Network (HBN) ** | MPRAGE | 0.8x0.8x0.8 | 320x320 | 224 | 1060 | 2500 | 130 | 2 | 7/8 (slice partial Fourier) off for phase | 8 | Prisma |
|  | MPRAGE+PMC | 1.0x1.0x1.0 | 256x256 | 176 | 1060 | 2500 | 240 | 2 | off | 8 | Prisma |

This ultrafast gradient-echo 3D pulse sequence is used by a large fraction of neuroimaging researchers because of its excellent contrast properties and capacity to collect reliable structural images for cortical thickness and volumetric measures (Wonderlick et al. 2009). MPRAGE can be considered as a defacto standard imaging sequence for brain morphometry studies^2^. As such, large neuroimaging studies such as the WU-Minn Human Connectome Project (HCP) (Van Essen et al. 2013) more recently referred to as the HCP Young-Adult (HCP-YA), the NKI-Rockland Sample (Nooner et al. 2012), the UK BioBank (Sudlow et al. 2015), and the Alzheimer's Disease Neuroimaging Initiative (ADNI) (Jack et al. 2008), all use the MPRAGE sequence to collect structural T1 weighted images of the brain. While some slight differences in sequence parameters exist across studies, such as voxel size and TR/TI values, differences are small across parameters (see Table 1
).

For navigator-based prospective motion correction (PMC) approaches, the sequence periodically collects fast-acquisition lower resolution images (navigators) to estimate the amount and direction of head motion since the last navigator was collected. Based on the motion estimation, sequence parameters are adjusted at each repetition time (TR) to nullify this motion. For MPRAGE sequences, navigators can be collected and motion can be estimated during the long inversion recovery time (TI) and applied to update the readout orientation for the current line of k-space. An early example of this method is the PROMO sequences (for GE scanners) that employ spiral acquisitions to collect navigators along the three cardinal planes of the volume (coronal, axial, and sagittal) (N. White et al. 2010; Sarlls et al. 2018). This has been extended by Tisdall et al. to use echo volume imaging (EPI applied to all 3 dimensions) to collect 3D vNAV (Tisdall et al. 2012, 2016). In addition to prospectively correcting for motion that occurs between acquisitions, this sequence with PMC can also identify large motion that occurs during an acquisition. The TRs that have motion above a predefined threshold are reacquired at the end of the sequence. The number of TRs that can be reacquired and motion threshold is set by the operator. The MPRAGE sequence with PMC (MPRAGE+PMC) has been widely adopted by research groups and more specifically, by new large imaging studies (see Table 1
). The equivalent of the vNav sequence for Philips MRIs is the iMOCO (Andersen et al. 2019).

Among those most attracted to the promises of sequences with PMC are pediatric imaging researchers (S. Y. Bookheimer 2000). In particular, head motion has been shown to significantly reduce gray matter volume and thickness estimates accuracy (Reuter et al. 2015) and also alter gray matter probability scores (Gilmore, Buser, and Hanson 2019). Given this concern, the longitudinal Adolescent Brain Cognitive Development (ABCD) Study (Casey et al. 2018) adopted the MPRAGE+PMC sequence as its standard T1-weighted structural sequence. The Healthy Brain Network (HBN) study (Alexander et al. 2017) also adopted the new MPRAGE+PMC sequence, while also maintaining the HCP style MPRAGE sequence due to concerns regarding reproducibility across sequences. The original HCP study (HCP-YA) is a project that has already concluded its data acquisition, but the study is now being expanded through the HCP Lifespan Studies. The Lifespan Studies have all adopted the MPRAGE+PMC protocol, in addition to standard MPRAGE. This includes the HCP Aging (HCP-A) (Susan Y. Bookheimer et al. 2019) for ages 36-100+ years old, the HCP Development (HCP-D) (Somerville et al. 2018) for ages 5-21 years old, and the Lifespan Baby Connectome Project (BCP) for children aged 0-5 years old (Howell et al. 2019). See Table 1
 for details regarding T1-weighted structural sequences used in large neuroimaging studies.

Given that new large imaging studies (i.e. HCP Lifespan and ABCD) are using the MPRAGE+PMC sequence to collect their structural data, we raise a key question: should other researchers switch from the well established MPRAGE sequence to the MPRAGE+PMC sequence? In this manuscript, we present quantifiable similarities and differences between the HCP style MPRAGE to the ABCD style MPRAGE+PMC sequence to address this question.

## Methods

### Neuroimaging Data

All neuroimaging data used in this study were collected as part of the Healthy Brain Network (HBM) Project (Alexander et al. 2017) and were acquired on a Siemens Prisma Fit with a 32 channel head coil located at the Citigroup Biomedical Imaging Center (CBIC) at Weill Cornell Medicine. A total of 465 imaging sessions were analyzed. Of these 465 participants, 348 completed the full HBN MRI protocol and are included in this study, with an age range of 5 to 21 years old (mean=11.3±3.6) which included 120 females and 228 males. The HBN protocol at CBIC includes two structural T1-weighted sequences, one based on the Human Connectome Project-YA (here referred to as the “MPRAGE” sequence), and another based on the ABCD study with the MPRAGE sequence with PMC (here referred to as the “MPRAGE+PMC” sequence). These sequences differ in other parameters as well, such as voxel and matrix size, bandwidth, and partial Fourier. Nonetheless, it is worth noting that the parameters for the MPRAGE and MPRAGE+PMC sequences were independently optimized by the designers of the HCP and ABCD studies (Glasser et al. 2013; Casey et al. 2018).

Another set of participants (N=72) performed a test-retest protocol (Test-Retest Group). As part of the protocol specifically designed for this study, these participants performed two MPRAGE scans and two vNav scans within the same imaging session. Specifically, one MPRAGE+PMC (MPRAGE+PMC1) and then one MPRAGE (MPRAGE1) sequence were performed at the beginning of the imaging session and the other two sequences were repeated at the end of the imaging session (MPRAGE2 and then MPRAGE+PMC2). This strategy was chosen since a larger amount of head motion is expected on the runs at the end of the session. The test-retest group had an age range of 5 to 20 years old (mean=11.6±3.7) with 23 females and 42 males. The HBN protocol and timing of the sequences for both groups are provided in Supplementary Tables 1 (ST1) and 2 (ST2).

### Imaging Parameters

The MR Protocol Guidance from the Human Connectome Project (Glasser et al. 2016) was followed to define the 3D MPRAGE HCP style imaging sequence. For the structural sequences with the navigators (MPRAGE+PMC), we used the protocol from the ABCD study (Casey et al. 2018). The imaging sequence protocol parameters used in this study are shown in Table 1
. Additionally, for the MPRAGE+PMC sequence, we configured the sequence with a reacquisition threshold of 0.5 (see equation [3] in (Tisdall et al. 2012) for details) and up to 24 TRs could be remeasured. The MPRAGE sequence had a duration of 7 minutes and 19 seconds, while the MPRAGE+PMC can take up to 7 minutes and 12 seconds to be acquired. During the structural runs, the participants were shown the Inscapes Movie (Vanderwal et al. 2015), a video developed to improve compliance related to motion and wakefulness.

### Visual Quality Control

An instance of the Braindr web application (Keshavan, Yeatman, and Rokem 2019) was created for this project to perform visual quality control of the MPRAGE and MPRAGE+PMC scans. This instance contained only the images collected for this study. Within Braindr, a rater can choose to “PASS” or “FAIL” an image depending on the quality. We asked the raters to cast their vote based on the general quality of the image but to specifically examine if the border areas between white and gray matter are blurry or not. Example images used for training are shown in Supplementary Figure SF1. Five research assistants of HBN participated as raters. They did not have any prior knowledge regarding the focus of this study (to compare MPRAGE and MPRAGE+PMC). For each structural image two axial and two sagittal slices were shown to the raters. Hence, every MRI image received a total of 20 votes. Slices were presented to the raters in random order.

### Quality Control

Six measures of quality control for the structural images were performed by using the Quality Assessment Protocol (QAP) toolbox (Zarrar et al. 2015). Specifically for each subject and structural image the following quality control scores measures were calculated:

- Contrast to Noise Ratio (CNR): measures the mean of the gray matter intensity values minus the mean of the white matter intensity values divided by the standard deviation of the values outside the brain) (Magnotta, Friedman, and FIRST BIRN 2006);
- Signal to Noise Ratio (SNR): measures the mean intensity within the gray matter divided by the standard deviation of the values outside the brain)(Magnotta, Friedman, and FIRST BIRN 2006);
- Foreground to Background Energy Ratio (FBER): measures the variance of voxels inside the brain divided by the variance of voxels outside the brain;
- Percent Artifact Voxels (PAV): measures the proportion of voxels outside the brain with artifacts to the total number of voxels outside the brain (Mortamet et al. 2009);
- Smoothness of Voxels (FWHM) measures the full-width half maximum of the spatial distribution of the image intensity values in voxel units (Magnotta, Friedman, and FIRST BIRN 2006);
- Entropy Focus Criterion (EFC), measures the Shannon entropy of voxel intensities proportional to the maximum possible entropy for a similarly sized image; Indicates ghosting and head motion-induced blurring) (Atkinson et al. 1997).

For the CNR, SNR, and FBER, a higher score means a better image, while for the FWHM, PAV, and EFC, a lower score is better.

Two additional quality control measurements were also performed that are not part of the QAP package. For the first additional measure, we compared the background noise in two regions around the brain. One 12mm radius circle located in front of the forehead immediately above the eyeball (Anterior ROI), and another above the head (Superior ROI) (See Supplementary Figure SF2 for the location of the circles). We then calculated the ratio of the average signal from the Anterior divided by the Superior region of interest, hence Anterior-to-Superior Ratio (ASR). The rationale of using ASR is that the prime source of image motion is manifested in the AP phase encode direction and results in increased noise and blurring in the spatial domain (Barish and Jara 1999), which we seek to quantify. Minimal motion is expected in the Inferior-Superior direction along the bore of the magnet. By comparing the background signal that is anterior to the head to a background signal superior to the head we are capturing the effects of head motion. A similar approach is performed by White et al. (White et al. 2018). A lower ASR yields lower anterior noise and hence, better image quality. The second additional quality control measure is Freesurfer’s Euler number, which summarizes the topological complexity of the reconstructed cortical surface and has been shown to identify “unusable” images with very high accuracy (Rosen et al. 2018). Higher Euler numbers were shown to be consistently positively correlated with manual image ratings.

### Structural Quantitative Measurements

We extracted morphometry measurements from the images using Mindboggle v1.2.2 (Klein et al. 2017). Within Mindboggle, Freesurfer v5.1 (Fischl 2012) measures were also extracted. These data were processed using an AWS EC2 r4.large instance with Amazon Linux AMI 2017.9 operating system. For each Freesurfer label, measurements included volume, area, median travel depth, geodesic depth, and the median measurement of Freesurfer’s cortical thickness, curvature, and convexity of the sulcus (Fischl 2012). Geodesic depth is the shortest distance along the surface of the brain from the point to where the brain surface makes contact with the outer reference surface (Klein et al. 2017), whereas travel depth is the shortest distance from a point to the outer reference surface without penetrating any surface (Giard et al. 2011; Klein et al. 2017). Total gray matter volume across different structural runs was also measured with FSL’s SIENAX (Smith et al. 2002) package, in addition to the results obtained through Mindboggle.

### Reliability

Intraclass correlation coefficients (ICC(3,1)) was used to calculate the reliability for each of the morphometric measures (Shrout and Fleiss 1979). Inter-sequence reliability was measured between the different imaging sequences, MPRAGE and MPRAGE+PMC. Intra-sequence reliability was measured across different imaging runs for the same pulse sequence.

### Age-Related Changes

We estimated age-related changes to compare the two imaging sequences. Age-related curves were separated by sex and by the quantity of motion during the functional sequences.

### Motion estimation

EPI volumetric navigators with an 8mm isotropic resolution are collected with the MPRAGE+PMC pulse sequence. These volumes are used as navigators to estimate the head motion during the scan, and one three dimensional volume is acquired at each TR. For the HBN study, the total number of volumes that were acquired at each MPRAGE+PMC sequence ranged from 143 to 168, depending on the number of TRs that need to be reacquired based on subject motion (Tisdall et al. 2012). By using QAP, which internally uses AFNI’s (Cox 2012) 3dvolreg to estimate motion parameters, the framewise displacement (FD) (Jenkinson et al. 2002) was calculated for each MPRAGE+PMC run on these low resolution EPI volumes^3^. The FD was then normalized to FD per minute (*FD_pm_*) by (Tisdall et al. 2012):

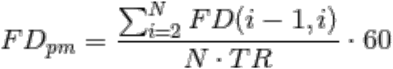

where *N* is the number of TRs, *TR* is the repetition time in seconds, *i* is the volume, and *FD(i-1,i)* is the FD between two subsequent volumes.

With the MPRAGE sequence, we cannot directly estimate motion, hence we investigated if the average motion across all functional scans can be used as a proxy for how much a participant moves during a structural scan (Pardoe, Kucharsky Hiess, and Kuzniecky 2016; Savalia et al. 2017). Specifically, the average *FD_pm_* for all functional MRI scans of the protocol were also calculated and compared.

## Results

### How do the Sequences Compare by Visual Inspection?

By visual inspection, there were some key differences in image intensity and quality when comparing the MPRAGE and MPRAGE+PMC sequences (Figure 1). For the purposes of demonstration, through visual inspection of the structural images, we identified two participants, one with a low amount of motion (Low-Mover) and one with a high amount of motion (Mover). For the participant data on the left (Low-Mover), visually, the images appear to be of excellent quality for both the MPRAGE and MPRAGE+PMC sequences. This subject had a low amount of motion during the data collection of the MPRAGE+PMC sequence (*FD_pm_* = 6.04 during MPRAGE+PMC sequence). For the functional scans of the protocol, the same participant has a low *FD_pm_* = 7.24 (see *Motion Estimation* section below on how head motion was estimated for the MPRAGE runs). The images on the right represent a participant with a large amount of motion for the MPRAGE and MPRAGE+PMC sequence. Even though there was a large amount of head motion during the acquisition in the MPRAGE+PMC sequence (*FD_pm_* = 62.23 during MPRAGE+PMC sequence; average *FD_pm_*=17.41 during functional scans), the quality of the T1’s is still sufficient for many applications. That is not the case for the MPRAGE images, where the ringing artifacts are strikingly pronounced and this data would have to be discarded for any neuroimaging study. However, it is important to note that for the MPRAGE+PMC image, the gray-white matter boundaries are not as sharp as the low motion subject. There are also some ringing artifacts present in the MPRAGE+PMC image. Hence, the MPRAGE+PMC image is not completely immune to head motion, as seen in Figure 1 and also in Supplementary Figure SF3.

**Figure 1.**
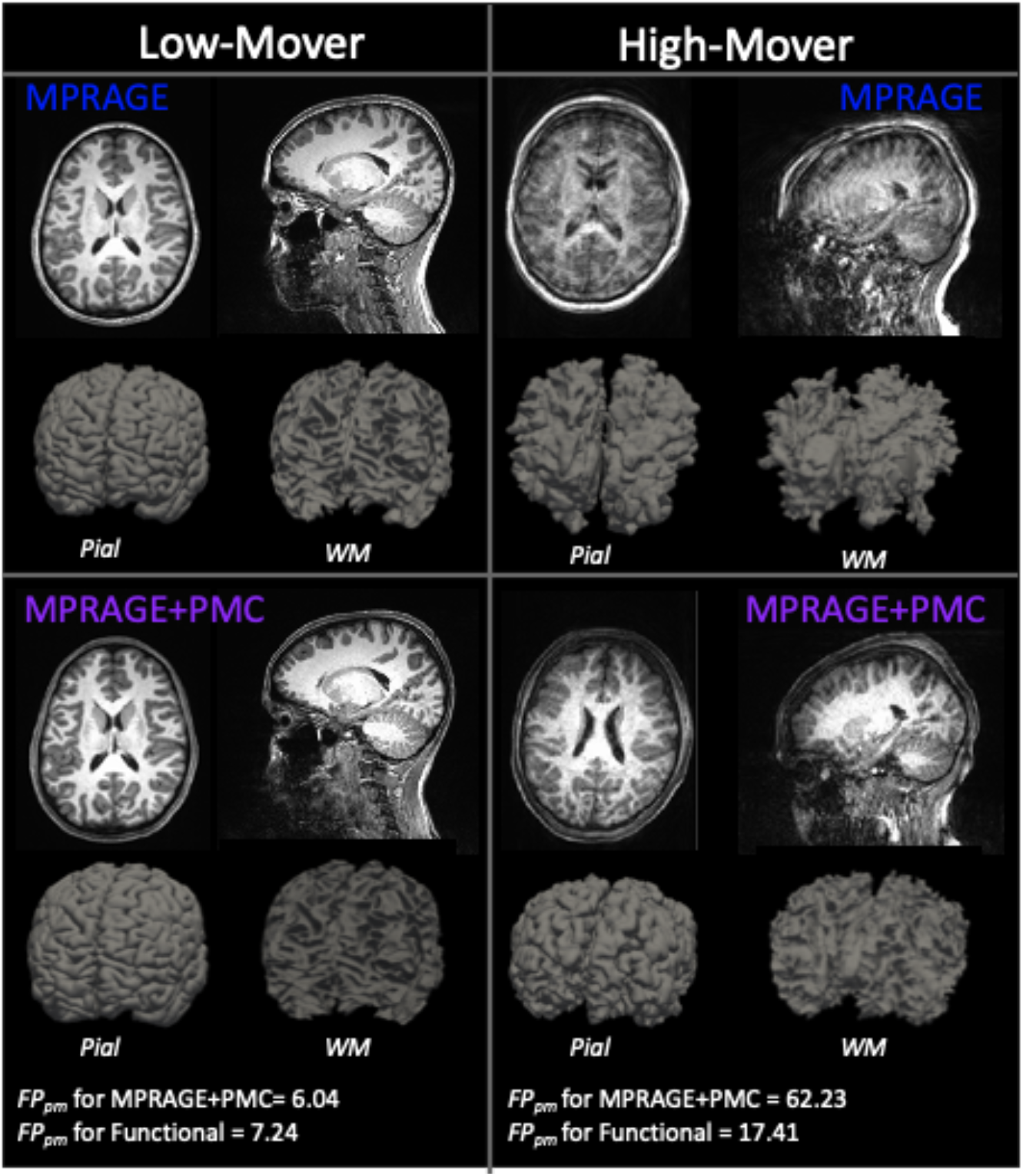
T1 structural images for the two sequences, MPRAGE and MPRAGE+PMC. The top row shows the MPRAGE sequence, while the bottom row shows the images that were generated with the MPRAGE+PMC sequence. Columns represent two different participants, one with minimal head motion (left, Low-Mover) and another with a large quantity of motion (right, High-Mover). Pial and white matter (WM) surface reconstruction from Freesurfer are also shown.

### Quantifying Visual Rating

By using Braindr, each of the 5 raters inspected a total of 3,920 slices. These images include scans from the large group, with a total of 346 participants that completed both the MPRAGE and MPRAGE+PMC sequences (346*2*4 = 2,768 slices). Images shown to the raters are also from the test-retest group, which includes 72 participants that completed the 4 structural runs, with the MPRAGE1 and MPRAGE+PMC1 sequences at the beginning of the session and MPRAGE2 and MPRAGE+PMC2 at the end. A total of 1,152 slices (72*4*4) from the test-retest group were shown to the raters.

Wilcoxon signed-rank tests were calculated to test if the observed differences are statistically significant. With the data from the large group, there was a significant (p<0.001) higher score for the MPRAGE+PMC images compared to the MPRAGE images with a z-score equal to −6.8298. Within the test-retest group, the MPRAGE-PMC scans were also significantly (p<0.001) scored higher than the MPRAGE scans, with a z-score = −5.0029 when comparing the two scans at the beginning of the run (MPRAGE1 and MPRAGE+PMC1) and a z-score = −4.9517 for the scans at the end of the run (MPRAGE2 and MPRAGE+PMC2). Likewise, we calculated the proportion of images from each scan type that had an average score above 0.5. For the large group, 60.06% and 72.99% of the MPRAGE and MPRAGE+PMC scans, respectively, have an average score above 0.5. For the test-retest group, there are 56.94% of the MPRAGE1, 44.44% of the MPRAGE2, 81.94% of the MPRAGE-PMC1, and 66.67% of the MPRAGE-PMC2 scans with a score above 0.5. Density plots of the average score per image type are shown in Figure 2 for the test-retest group.

**Figure 2.**
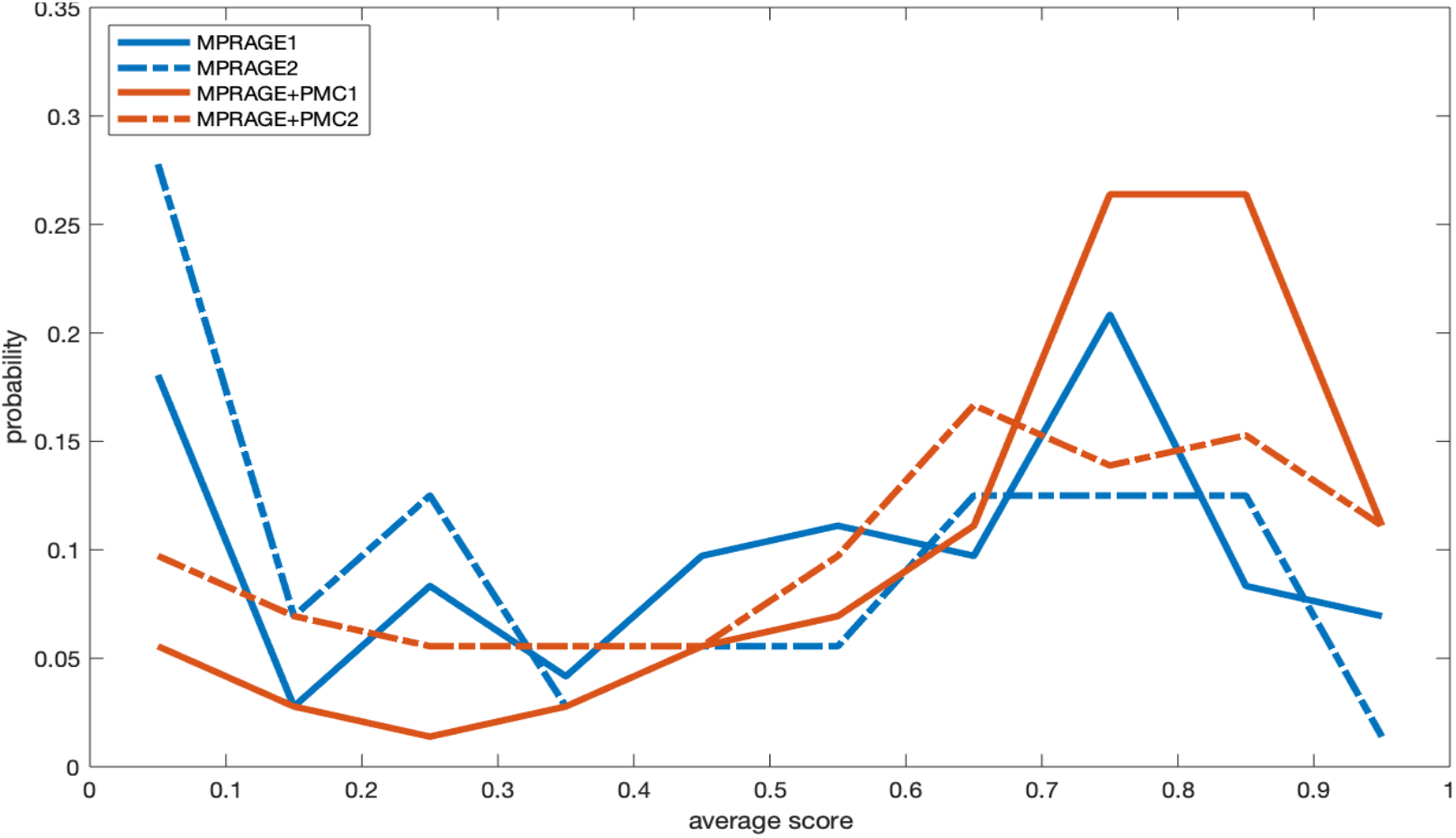
Density plots of the average Braindr score of each scan type for the test-retest group.

### Does the Surface Reconstruction Complete?

Another simple, however very practical, comparison of these two sequences is to test whether Mindboggle was able to complete processing the surface reconstruction. With images of poor quality, such as the one seen for the “High-Mover” subject in the MPRAGE sequence (Figure 1), the software does not complete and returns an error instead of the morphometry measurements. For all participants in which there was an error, incomplete surface reconstructions were caused by the topological defects. This was either due to the failure of the automatic topology fixer (mris_fix_topology) or as a result of the running time exceeding the time limit (60 hours) that was set. Considering that the purpose of this study is to show the differences and similarities between the MPRAGE and MPRAGE+PMC sequences, we did not manually edit and correct the volume and attempt to reprocess the data as is suggested in FreeSurfer’s tutorials.

For the large group, a total of 346 participants completed both the MPRAGE and MPRAGE+PMC sequences. Of the 346 images, Mindboggle successfully completed surface reconstruction in 89.3% (N=309) of the MPRAGE images and 91.6% (N=317) of the MPRAGE+PMC images. Using McNemar’s test (McNEMAR 1947), no statistically significant differences (p>0.1) were found between completing the processing of the images for the different sequences. We visually inspected the 31 images from the MPRAGE+PMC images that did not complete surface reconstruction and all looked blurry (see examples in Supplementary Figure SF3). The average motion from the EPI navigators of the MPRAGE+PMC sequences was then calculated. For the images that completed the surface reconstruction, there was an *FD_pm_* = 13.70 +- 12.85, while for the images that were not completed the motion was much higher, with an *FD_pm_* = 47.56 +- 37.32. For the subsequent analysis shown in this manuscript that depends on the surface reconstruction results, only the data from participants that Mindboggle was able to complete processing both images are used. Hence, the following results that are presented only use data that has already passed through a first level of quality control, i.e. completing surface reconstruction. A total of 290 subjects (105 females, mean age = 11.20 +- 3.66, age range = [5.44, 20.47]) are included in the following analyses that depend on surface reconstruction estimates.

Of the 72 test-retest participants that completed all four runs, Mindboggle completed processing on 94.5% (N=68) of the MPRAGE1, 98.6% (N=71) of the MPRAGE+PMC1, 97.2.6% (N=70) of the MPRAGE2, and 97.2% (N=70) of the MPRAGE+PMC2. Again, using McNemar’s test we found no statistically significant differences (p>0.1) between the sequences regarding Mindboggle completing the processing of the images. Mindboggle was able to calculate morphometric measurements in all 4 structural runs for 65 participants. These participants are used in the reliability tests shown below.

Even though there was no significant difference in surface reconstruction completion between MPRAGE and MPRAGE+PMC scans, it does not indicate the two sequences have the same data quality or accuracy measurements. As the reliability results are shown below, there are significant differences in reliability between the scans in which the surface reconstruction completed.

### Reliability

Intra- and inter-sequence reliability results for the Mindboggle measurements are shown in Figure 3. ICC scores are shown for each of 62 cortical regions from the Desikan Atlas (Desikan et al. 2006), which are sorted by Yeo networks (Yeo et al. 2011; A. F. Alexander-Bloch et al. 2018) (list of regions can be seen in Supplementary Table ST3), and for the following measurements, (1) area, (2) Freesurfer median cortical thickness, (3) travel depth, (4) geodesic distance, (5) curvature, and (6) convexity. The first row shows results for all participants. They were then divided into two groups, through a median split of the mean *FD_pm_* of the functional scans. The Intra-sequence reliability of cortical measures extracted from the two MPRAGE+PMC sequences was significantly higher (paired t-test, all p < 0.0001) for all of the regions and measures compared to the MPRAGE pair without PMC. These significant results are observed when including all subjects in the analysis, and also when calculating the t-tests with only the low- or high-motion subjects. These results are possibly due to a higher motion during the MPRAGE2 scan, which was collected at the end of the session. Another noticeable result is that, even for the low motion subjects, the ICC was significantly higher (paired t-test, all p < 0.0001) for the MPRAGE1 × MPRAGE+PMC1 pair (inter-sequence) compared to the MPRAGE1 × MPRAGE2 pair (intra-sequence). Again, this is possibly a result of the higher motion in the MPRAGE2 runs, even though these were the subjects with lower motion estimation scores. From the same reproducibility results it is important to notice that even though the voxel sizes are of different sizes for the MPRAGE (0.512mm^3^) and MPRAGE+PMC (1mm^3^) sequences, the ICC scores between MPRAGE1 and MPRAGE+PMC1 are higher than repeating the MPRAGE sequence.

**Figure 3.**
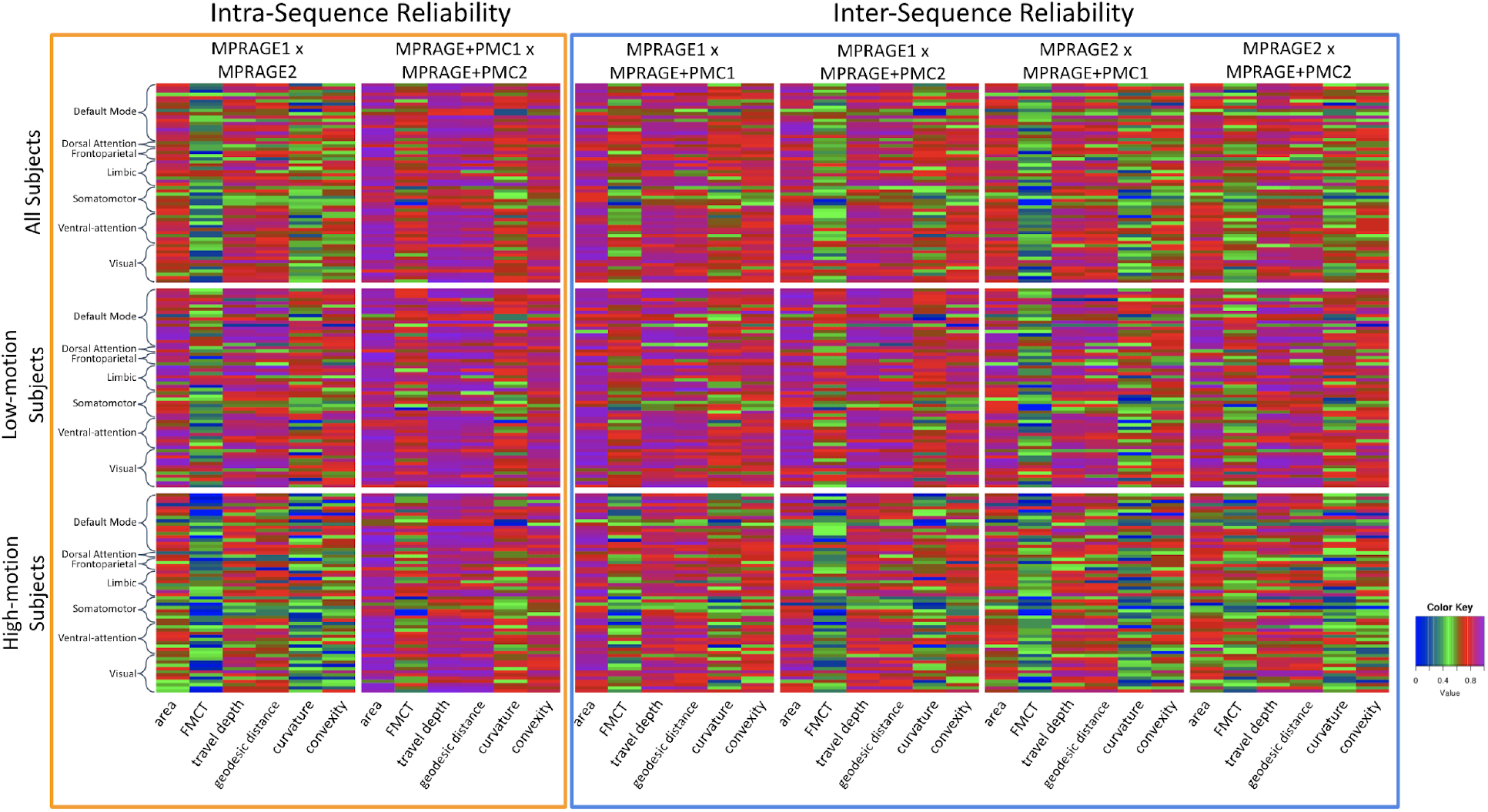
Test-retest reliabilityICC results for Mindboggle measurements within each of the 62 Desikan-Killiany Atlas cortical regions. Regions have been sorted by the Yeo 7 Network Atlas (Yeo et al. 2011). Measurements tested were (1) area, (2) Freesurfer median cortical thickness (FMCT), (3) travel depth, (4) geodesic distance, (5) curvature, and (6) convexity.

Density plots of ICC for the area, volume, and Freesurfer median cortical thickness across all brain regions are shown in Figure 4. As can be seen in the density plots, the pair MPRAGE+PMC1 × MPRAGE+PMC2 (orange line) outperformed the ICC scores of all other pairs. It is clear that the reproducibility between MPRAGE+PMC1 and MPRAGE+PMC2 is high, with an average ICC score above 0.8 for Area and Volume and above 0.6 for cortical thickness. The pair MPRAGE1 × MPRAGE+PMC1 (green line) typically showed the second-best performance. The lower ICC scores in the low-motion and high-motion subjects for any pair that contains the MPRAGE2 run (blue, purple, and brown lines) are highly observable, especially for Area. The MPRAGE+PMC2 run is also performed at the end of the session, however contrary to the MPRAGE sequence, we can directly measure the amount of motion during that run. During the MPRAGE+PMC2 run, on average, there is at least twice the amount of head motion compared to MPRAGE+PMC1 (*FD_pm_* = 7.5 for MPRAGE+PMC1 and *FD_pm_* = 15.34 for MPRAGE+PMC2). Also when considering the pairs that contain the MPRAGE2 run, there is a large negative shift in ICC scores when comparing the “Low-Motion” and “High-Motion” subjects. With the other pairs, there is also a negative shift in ICC scores, but at a much smaller scale. These results corroborate with the notion that the MPRAGE+PMC is more robust to motion compared to the sequence without PMC.

**Figure 4.**
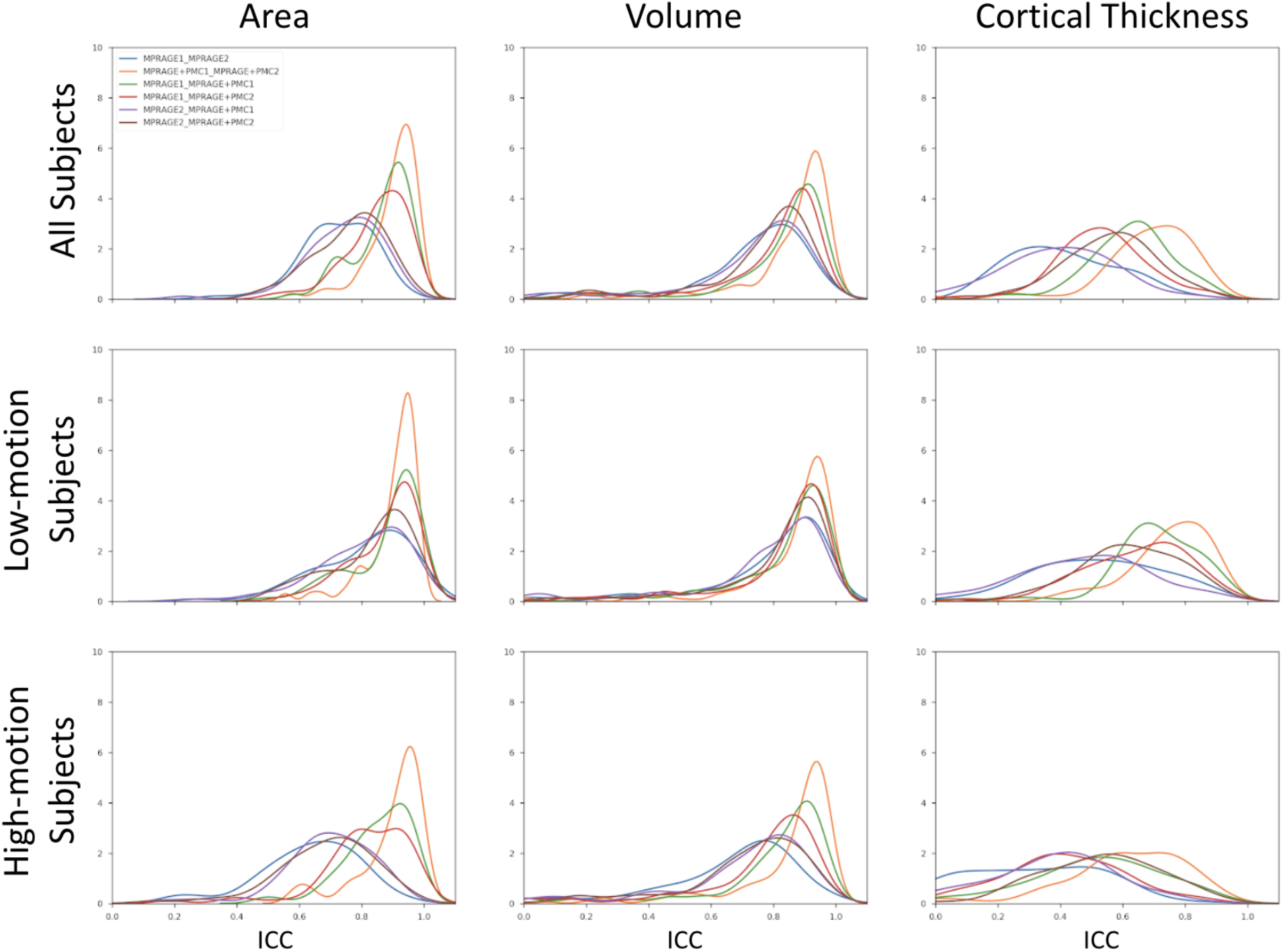
Density plots of ICC for the test-retest group that performed two MPRAGE scans and two MPRAGE+PMC scans within the same session. ICC is calculated for Area, Volume, and Cortical Thickness.

ICC of Cortical Thickness was worse for all pairs compared to Area and Volume. This shows how sensitive the measurement of Cortical Thickness is, especially in regards to head motion. The improvement in ICC for the MPRAGE+PMC pair over the MPRAGE pair was unanticipated for the low motion group, given that the MPRAGE sequence has a better spatial resolution, which is expected to obtain better cortical thickness estimation results. Another key result from the “ideal” low motion group, is that the inter-sequence pairs MPRAGE1 - MPRAGE+PMC1 (green line) and MPRAGE1 - MPRAGE+PMC2 (red line) show higher reliability than the intra-sequence pair MPRAGE1-2 (blue line) for all the three measures being evaluated.

Besides the reliability of measures from all 62 individual brain regions, the impact of acquisition sequence on the overall gray matter volume estimation was evaluated using Mindboggle and SIENAX (Figure 5) with the test-retest dataset. Pairwise comparisons were made for each combination by calculating the absolute difference in volume measures. MPRAGE+PMC1 and MPRAGE+PMC2 had the most similar gray matter volumes for both toolboxes. The largest differences were observed in the pairs that included the MPRAGE2 image. These results indicate that the prospective motion correction sequence is robust for measuring gray matter volume regardless of the toolbox used to calculate volumes. It also endorses the assumption that the MPRAGE+PMC sequence provides us more reliable results compared to MPRAGE, independent of when the structural sequence is performed within the session, beginning or end. Outliers are cases where the brain extraction failed in at least one of the images, mostly caused by high head motion.

**Figure 5.**
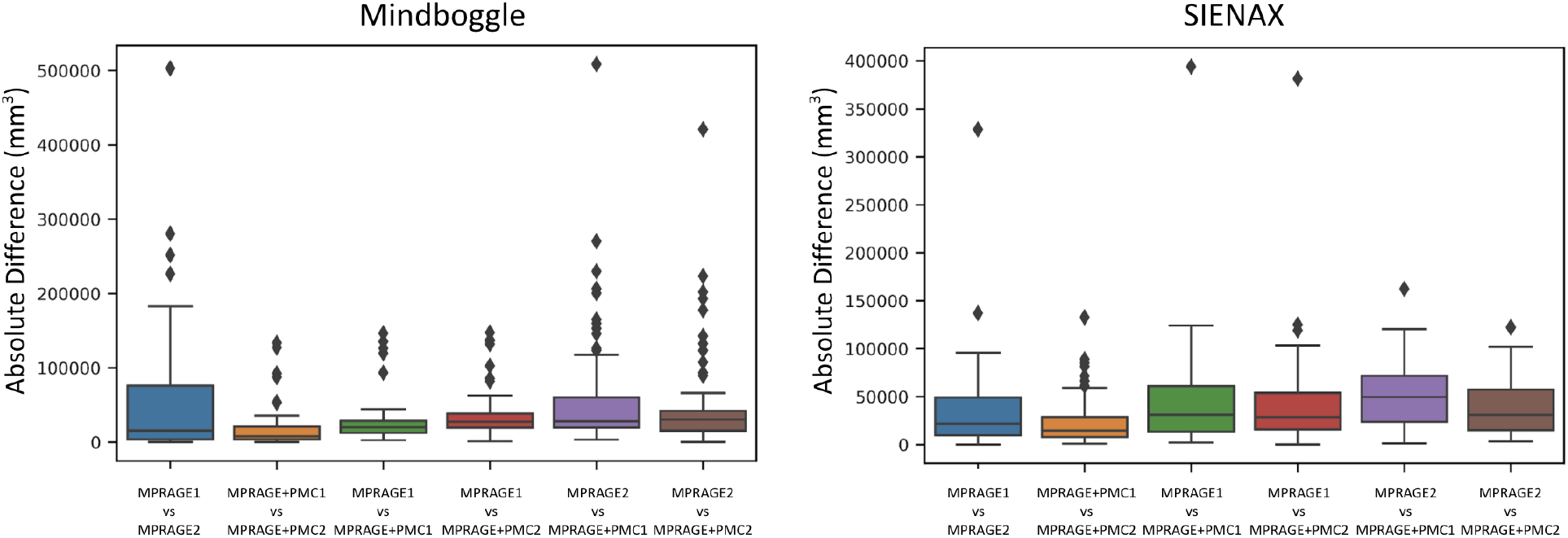
The absolute difference in gray matter volume within the test-retest group. Gray matter was measured using MindBoggle and SIENAX.

### How does Motion Affect Structural Measurements Between Sequences?

With the larger dataset, we performed an analysis to investigate if the differences in measurements of cortical thickness are affected by head motion. For each region, a partial correlation was calculated between the difference in cortical thickness measured with the MPRAGE and MPRAGE+PMC images (MPRAGE - MPRAGE+PMC) and the mean *FD_pm_* across the functional scans, controlling for age and sex. Of the 62 regions in the atlas, 26 showed a significant negative correlation (p<0.05) between the difference in cortical thickness in the images and the motion estimation, mostly located in the frontal, parietal and temporal lobes. Only one region showed a significant positive correlation, the right Isthmus of the cingulate cortex. These results indicate that, as there was an increase in subject head motion the difference in the measurement of cortical thickness between MPRAGE and MPRAGE+PMC increases, with a larger cortical thickness estimate in the MPRAGE+PMC sequence in 26 regions (see Figure 6A and Supplementary Table ST3 for the partial correlation scores for all regions).

**Figure 6 -.**
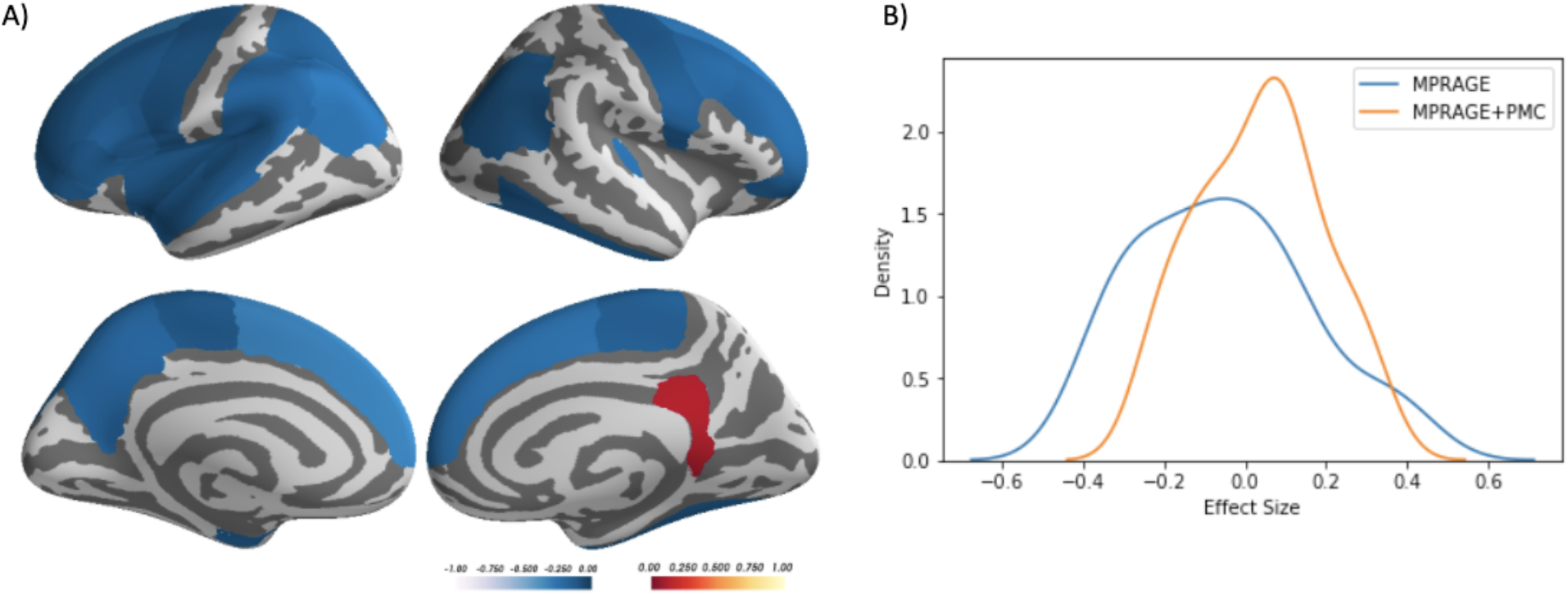
A) Desikan Atlas regions that showed a significant partial correlation (p<0.05), corrected by age and sex, between the difference in cortical thickness measurements (MPRAGE-MPRAGE+PMC) and mean FD across the functional scans. B) The distribution of the effect size (correlation of cortical thickness and motion estimates) across cortical regions for each sequence.

To evaluate motion related bias, the effect size (correlation) was calculated between the estimated cortical thickness values and the mean FD_pm_ for each sequence separately. The distribution of the effect size across cortical regions is shown in Figure 6B. A paired t-test shows that the effect size across all brain regions are significantly different between the MPRAGE and MPRAGE+PMC sequences (p < 0.0001, MPRAGE mean = −0.055, std = 0.211; MPRAGE+PMC mean = 0.036, std = 0.152).

We also calculated pairwise t-tests to compare the cortical thickness measurements between MPRAGE and MPRAGE+PMC in the 62 cortical regions from the Desikan Atlas. The t-tests were controlled for age, sex, and mean FD for the functional scans. Results showed that there was a significant difference (p<0.05) in 13 of the 62 regions. Of these 13, 11 showed a larger cortical thickness for MPRAGE and 2 for MPRAGE+PMC (See Supplementary Figure SF4). Statistical scores for all the regions are shown in Supplementary Table ST4.

### Age-Related Differences

Figure 7 shows development curves for total volume, gray and white matter volume, and ventricle volume for male and female participants. Only a smaller subset of subjects (N=248, 92 females) was used to calculate the development curves since there were very few subjects older than 16 to obtain adequate development estimation curves at higher ages. Hence, development curves are shown only for ages 6 to 16. Black dots represent volumes calculated with the MPRAGE sequence while red dots represent the MPRAGE+PMC sequence. For each sequence, a quadratic curve was fit for estimating development growth. For the “All Subjects” the development graphs appear to be similar for both imaging sequences. With a median split, participants were grouped by low and high motion. Even for the subjects with large motion, the development curves also appear to be similar, just deviating at the higher ages for both groups. This deviation in curvature is possibly caused by the low amount of subjects that are older with a higher amount of motion. The development curves are shown for cortical thickness measurements for males and females are shown in Supplementary Figures SF5 and SF6, respectively.

**Figure 7.**
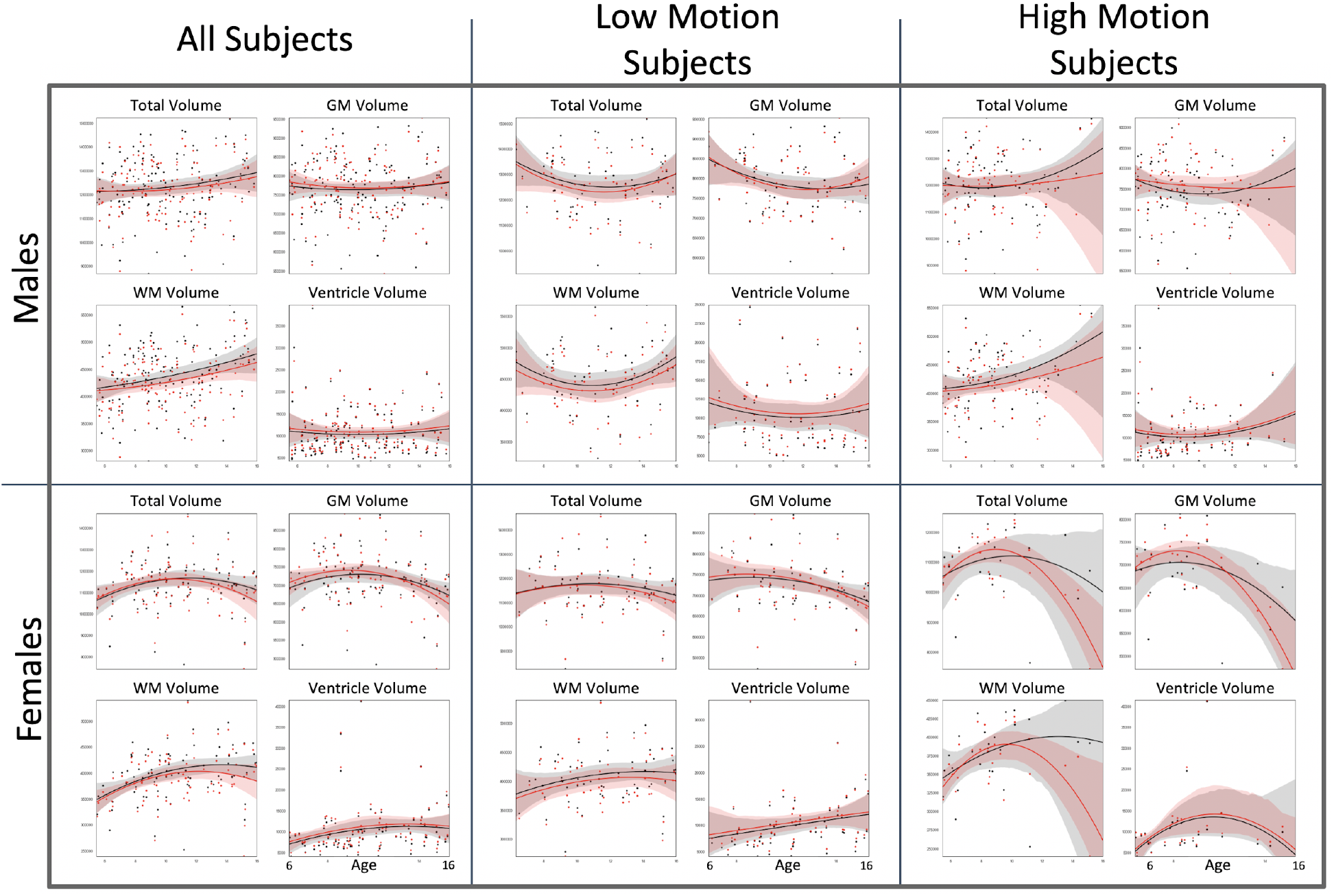
Black dots and lines (with 95% confidence intervals) are developmental measures for the MPRAGE sequence, while the red dots and lines are for the MPRAGE+PMC sequence. GM: Gray Matter; WM: White Matter.

To evaluate the effect of sequence type on volume measurement, we constructed 24 ANCOVAs, with age and mean *FD_pm_* across the functional runs as covariates (Savalia et al. 2017). Volume measurement (total brain, gray matter volume, white matter, and ventricle volume) was defined as the dependent variable and the sequence type (MPRAGE or MPRAGE+PMC) as the between factor. Including all the male participants in the statistical analysis, a statistically significant difference between the sequences was only found for white matter volume (F(2,675)=3.083, uncorrected P=0.046). For females, the only significant difference between sequences was found for ventricle volume (F(2,378)=3.061, uncorrected P=0.028). Within the low motion subjects, there are no statistically significant differences (p<0.05) between sequences for all volume measurements and both sexes. For the high motion group, the only significant result found was for ventricle volume in females(F(3,138)=2.771, uncorrected P=0.044). However, if we were to apply the Bonferroni correction for multiple comparisons, these significant findings would then be considered non-significant.

### Quality Control Metrics

The MPRAGE+PMC pulse sequence is identical to the MPRAGE sequence, except for the inclusion of a navigator acquisition and registration block, lasting 355 milliseconds, during the inversion recovery time and just before the parent sequence’s readout (Tisdall et al. 2012). In previous comparisons between these sequences, the MPRAGE+PMC sequence resulted in an approximately 1% reduction in contrast and a 3% reduction in image intensities (Tisdall et al. 2012). Importantly, these reductions were spatially uniform, so did not increase regional variation in image intensity (e.g., the ‘bias’ field). The MPRAGE+PMC sequence has been shown to result in more artifacts, such as ghosting, in the background, but since they did not overlap with the brain, they were not considered problematic (Tisdall et al. 2012).

In our study, acquisition parameters were identical between sequences, with the exception of voxel resolution, bandwidth, and partial Fourier. These differences are a consequence of MPRAGE+PMC’s navigate and register block reducing the amount of time available for parent sequence readout. From MRI theory, SNR is proportional to voxel volume (V) and inversely proportional to the square root of bandwidth (BW) and the square root of the partial Fourier reduction factor (R). The relative SNR from the MPRAGE+PMC sequence to the MPRAGE sequence can be calculated by:

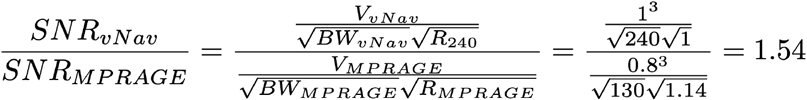

From these relationships, the SNR of the MPRAGE+PMC sequence is expected to be about 1.54 times greater than the SNR of the MPRAGE sequence (Craddock et al. 2013; Brown et al. 2014).

Figure 8 and shows the quality control metrics for the 348 participants. Paired t-tests were calculated at each measure to statistically compare sequences. Results showed significant differences (p<0.05) for all measures. Statistical scores are included in Figure 8. When comparing MPRAGE and MPRAGE+PMC, MPRAGE+PMC had a better score for CNR, ASR, and Euler number. MPRAGE exhibited a better score in all the other measures. Plots showing the quality control metrics separated by low and high-movers and also for the test-retest group are shown in Supplementary Figures SF7 and SF8, respectively.

**Figure 8.**
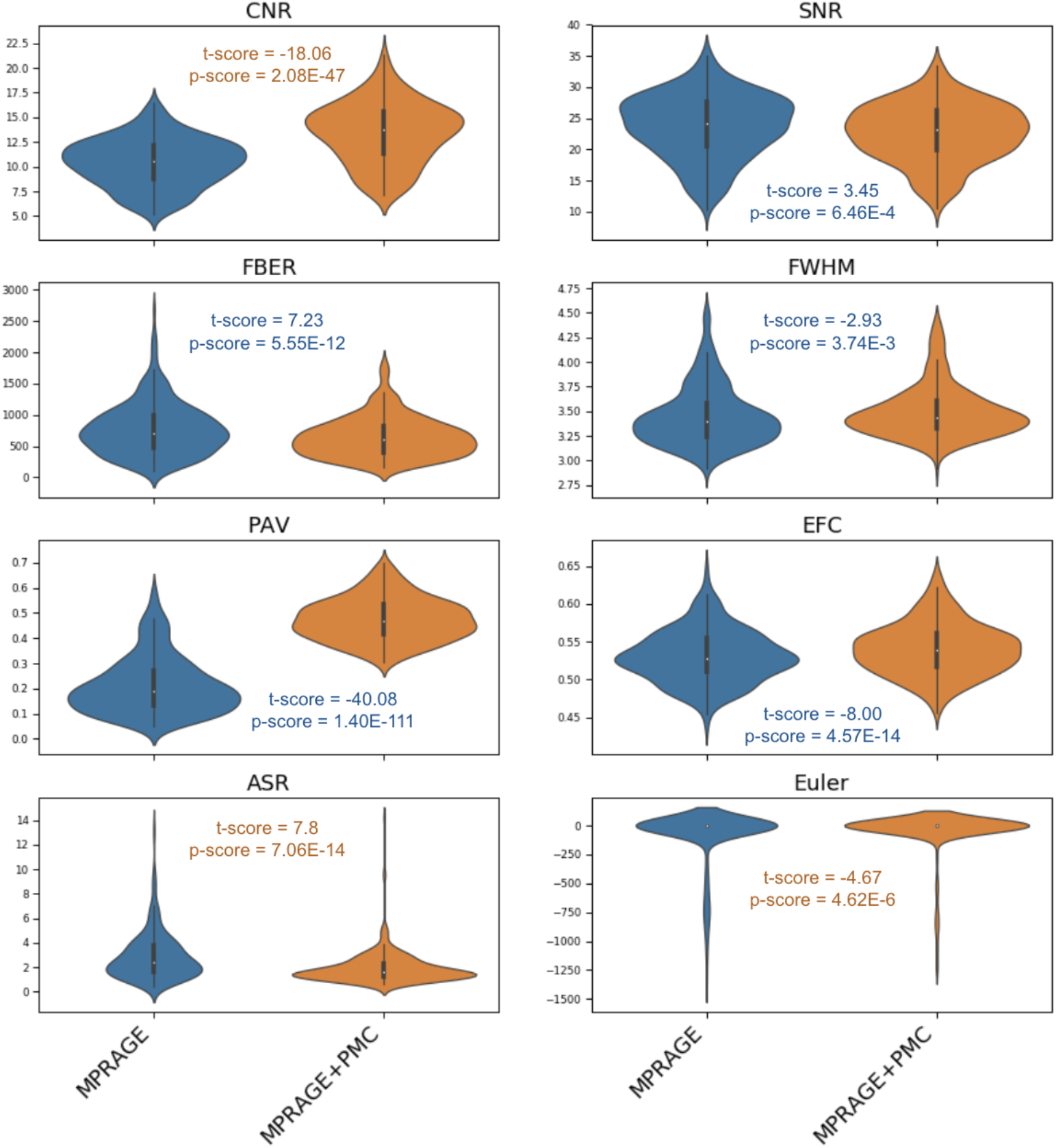
Quality control metrics for the MPRAGE and MPRAGE+PMC images across 287 participants. Metrics include; Contrast to Noise Ratio (CNR), Signal to Noise Ratio (SNR), (FBER), Smoothness of Voxels (FWHM), Percent Artifact Voxels (PAV), Entropy Focus Criterion (EFC), Anterior-to-Superior Ratio (ASR), and Freesurfer’s Euler number. Results of the paired t-tests comparing each of the quality control metrics are also shown. The t-scores and p-values are color-coded to indicate which image (MPRAGE or MPRAGE+PMC) performed better at each paired comparison, blue for MPRAGE and orange for MPRAGE+PMC.

These results are a bit unexpected, especially for SNR, since theoretically, the MPRAGE+PMC sequence should have an SNR 1.54 times greater than MPRAGE. Tisdal et. al (Tisdall et al. 2012) reported that the MPRAGE+PMC images had an increased ghosting effect that was only observed in the background. We are calculating SNR of each image by measuring the mean intensity within the gray matter and dividing by the standard deviation of the voxels outside of the brain. The increase in ghosting artifacts in the background would justify the reduction in SNR for the MPRAGE+PMC images. The same holds for justifying the inferior scores for MPRAGE+PMC in FBER, PAV, and EFC, which all depend on the background signal to calculate their metrics. The larger receive bandwidth (RBW) of the vNav sequence (240 kHz) compared to the RBW of the MPRAGE (130 kHz) might justify the increase in background noise, since larger RBW lets in more noise in the echo. The lower FWHM scores for the MPRAGE image are due to smaller voxel sizes compared to the MPRAGE+PMC image.

### Motion Estimation

Head motion occurring during fMRI scans has been proposed as a surrogate for sMRI motion when no other method for estimating motion from the data exists (Pardoe, Kucharsky Hiess, and Kuzniecky 2016; Savalia et al. 2017). The potential accuracy of fMRI as a surrogate of sMRI motion is supported by observations of high test-retest reliability for motion parameters across scans and sessions (Yan et al. 2013). But, fatigue, discomfort, and other factors are known to increase motion over time, which will likely degrade the surrogate’s accuracy. We directly tested the validity of using fMRI motion as a surrogate for sMRI motion by correlating sMRI motion estimates from the MPRAGE+PMC sequence with the motion from each of the fMRI scans collected in the same session with the test-retest dataset (see Supplementary Tables ST1 and ST2 and Alexander (Alexander et al. 2017) for details on the full imaging session.). We additionally tested how well the average motion across all fMRI scans correlates with the motion calculated in the MPRAGE+PMC run (Figure 9). In Figure 9 runs are listed in the order that they were collected.

**Figure 9.**
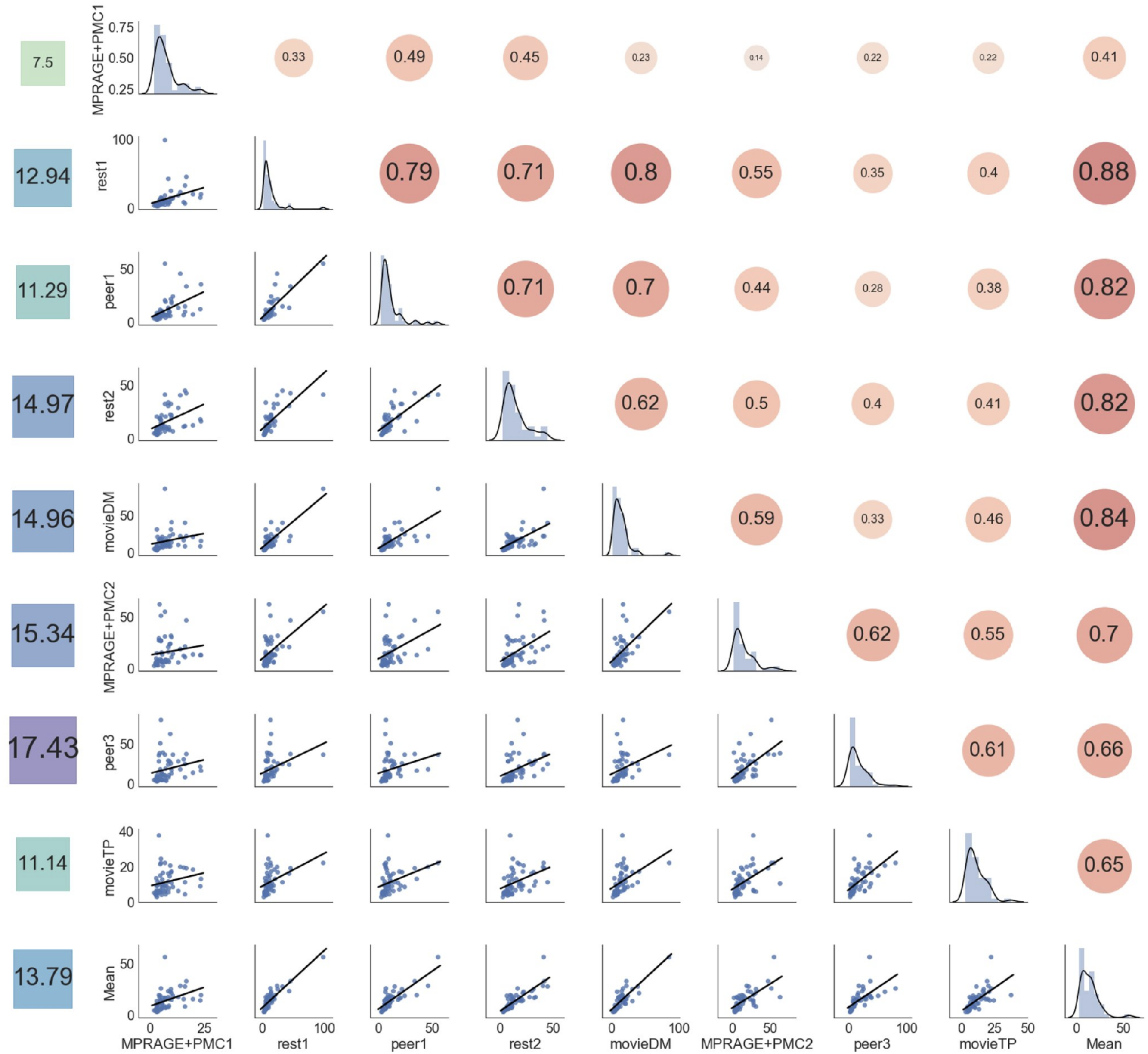
Correlation of the *FD_pm_* of all the runs. The main diagonal shows the distribution of *FD_pm_* each run (MPRAGE+PMC1 to movieTP) and the average *FD_pm_* of the functional runs (“Mean”). The bottom left of the diagonal shows the scatter plot of the motion parameters across runs, while the top right of the diagonal. The values on the left column show the average *FD_pm_* for each run.

The leftmost column of Figure 9 shows the average *FD_pm_*. As expected, there is a tendency for an increase in head motion as the session prolonged. There were two runs that were repeated in the first half and in the second half of the session, MPRAGE+PMC1-2 and peer1-3^4^. For the vMPRAGE+PMC runs, there was an increase in average *FD_pm_* from 7.50 to 15.34 for MPRAGE+PMC1 and MPRAGE+PMC2 respectively. A large increase in motion also occurs for the peer runs, from 11.29 in peer1 to 17.43 in peer3. An exception to this increase in head motion was MovieTP, which is a short animated and engaging movie (“The Present”), which could have been the reason for lower head motion during that run.

Results indicate that runs that were collected close in time showed higher correlations than runs that were collected farther apart. For example, rest1 has a much higher correlation with peer1 in *FD_pm_* (r=0.79) which is collected immediately after rest1, than it did to movieTP (r=0.40) which occurred at the very end of the scanning session. The exception to this order effect was rest1 and movieDM, which were separated in time, with a correlation of r=0.80. The average *FD_pm_* across the functional runs (“Mean”) exhibited a high correlation with all the runs, with values ranging from r=[0.41, 0.83]. The correlation of *FD_pm_* across the functional runs with the two structural runs was r=0.41 and r=0.70 for MPRAGE+PMC1 and MPRAGE+PMC2, respectively. Therefore, as previously suggested, the mean FD across the functional scans is a decent surrogate measure for the head motion for the structural scans (Pardoe, Kucharsky Hiess, and Kuzniecky 2016; Savalia et al. 2017).

## Discussion

The present study examined the relative advantages and interchangeability of the traditional MPRAGE and MPRAGE+PMC pulse sequences. Intra-sequence reliability demonstrated a clear advantage for the MPRAGE+PMC sequence in a hyperkinetic population, largely owing to the compromises in MPRAGE reliability among the higher movers. Inter-sequence reliabilities among low-motion participants demonstrated high comparability for the assessment of individual differences, suggesting the potential to change sequences mid-study when possible. In comparison to other studies that directly contrast the MPRAGE and MPRAGE+PMC sequences in a controlled environment, we tested the MPRAGE+PMC sequence in a “real world” scenario, where we did not explicitly ask subjects to move or maintain still during the acquisition of the structural images (Tisdall et al. 2016; Sarlls et al. 2018; Andersen et al. 2019). All subjects were requested to maintain their head as still as possible throughout the imaging session.

### Advantages and Disadvantages

Intra-sequence reliability scores of the two sequences revealed a clear superiority of the newer pulse sequence (MPRAGE-PMC). This was demonstrated across the broad range of morphometric measurements tested. The higher robustness to head motion observed for the MPRAGE-PMC sequence is not only owing to the adaptation of the gradients to motion, but the acquisition of TRs with large displacement (Tisdall et al. 2012), which is a similar strategy that was previously developed for the PROMO sequence for the GE platform (N. White et al. 2010).

It is worth noting that not all measures favored the sequence with PMC. As observed with the quality control indexes, there is a decrease in signal quality in the MPRAGE+PMC compared to MPRAGE. The MPRAGE sequence is superior to the MPRAGE+PMC sequence in 5 out of the 8 quality control measurements. Nonetheless, most of the measures in which the MPRAGE sequence is superior depends on the level of noise in the background, which in most part do not affect brain segmentation algorithms (i.e. Freesurfer, Mindboggle, Siena). The Euler number, in which MPRAGE+PMC is superior, actually can be considered a crucial quality control metric since it has been shown to directly correlate with manual ratings (Rosen et al. 2018). Additionally, the CNR is superior for the MPRAGE+PMC sequence. CNR is also a central quality control metric since it measures the contrast between the gray matter and the white matter intensities. This contrast is necessary for accurately finding the gray matter - white matter boundary, which is imperative for performing segmentation of brain areas/volumes and measuring cortical thickness.

### Should researchers switch sequences?

Inter-sequence reliability scores showed excellent mean ICC scores (>0.8) for the majority of the morphometric measures tested. The only exception is cortical thickness, which showed a mean ICC score of 0.512 between MPRAGE1 and MPRAGE+PMC1. Nonetheless, our results show higher inter-sequence reliability (MPRAGE1 × MPRAGE+PMC1) than intra-sequence reliability of the more traditional sequence (MPRAGE1 × MPRAGE2) in all the morphometric measures. As expected, inter- and intra-sequence reliability is higher for lower motion subjects compared to the higher motion subjects. Analogous results between the two sequences were also obtained for the development curves. This further corroborates with the notion that brain quantitative measures obtained in the different sequences are more equivalent then different. Additionally, the results obtained through visual inspection (using Braindr) showed that the images generated by the MPRAGE+PMC sequence were preferred by the raters.

Taking into account all considerations, we recommend that researchers: 1) use MPRAGE+PMC as their structural T1 weighted pulse imaging sequence for future studies, and 2) consider switching to the MPRAGE+PMC for ongoing studies. The first recommendation is relatively obvious given our findings that there are higher intra-sequence reliability scores for the MPRAGE+PMC pair (MPRAGE+PMC1 × MPRAGE+PMC1) compared to all other pairs of sequences. In contrast, the second recommendation may be somewhat surprising to some; however, it reflects our findings of high inter-sequence reliability, especially in the MPRAGE1 × MPRAGE-PMC1 pair. The immediate switch to MPRAGE+PMC sequence is likely most important to studies dealing with hyperkinetic populations, where findings are increasingly being questioned in view of associations with head motion (Reuter et al. 2015; A. Alexander-Bloch et al. 2016; Pardoe, Kucharsky Hiess, and Kuzniecky 2016). Neuroimaging researchers with projects studying low head motion participants may very well consider staying with the MPRAGE sequence, possibly finding some advantage given the higher quality control measures.

Beyond the quality control metrics, the only downside that we can identify for adopting the MPRAGE-PMC sequence is the potential increase in acquisition time. However, this increase in acquisition time is mostly caused by the repetition of TRs that surpass a motion threshold. If you are studying a population with high motion, on average this is actually an overall reduction in scan time, especially if researchers are considering repeating a full acquisition for high movers. The HBN initiative with participants aged 6-21 adopted a maximum repeat of 24 TRs. If time is of the essence and the study is with a low moving population, the maximum number of repeated TRs can be reduced to save scanner time.

It is important to note that the MPRAGE+PMC sequence is not entirely immune to head motion and other measures to restrain motion should be used in conjunction with this new sequence. Fortunately, the PMC pulse sequences can be used in conjunction with other strategies for minimizing head movements, such as training the subject in a mock scanner to get acclimated to the environment (de Bie et al. 2010), movie watching to reduce motion (Vanderwal et al. 2015; Greene et al. 2018), and other methods such as using customized head restraints have also been proposed (Power et al. 2019). Additionally, methods that quickly quantify the quality of the structural images have also been proposed (T. White et al. 2018), hence if necessary, a structural scan can quickly be repeated within the same session.

### Limitations

This study is limited in the sense that we do not have a direct measurement for motion during the MPRAGE sequence. However, we have attempted to estimate the motion by using the average motion across the functional runs. We have also not performed any rigorous visual inspection (Iscan et al. 2015) or post-processing quality control on the morphometric measurements (Ducharme et al. 2016). We did not want to discard any data due to poor image quality through visual inspection, hence directly comparing the two sequences. As can be seen in Figure 1, though a visual inspection for the high motion participant, we would probably discard the MPRAGE image but not the MPRAGE+PMC image. Another limitation of this study is that the voxel size of the sequences that we are testing are of different sizes and that the Bandwidth and Partial Fourier values differ between sequences. However, these parameters were independently optimized by the HCP and ABCD groups for the MPRAGE and MPRAGE+PMC sequences, respectively. Previous work by Tisdal et. al (2016) has directly compared the MPRAGE sequence with and without PMC using the same parameters. Attempting to find parameters that would be acceptable for both the MPRAGE and MPRAGE+PMC sequences is beyond the scope of this project, and would also entail that we would be using suboptimal parameters for both sequences. The objective of this study is to compare two T1-weighted MRI sequences that are used by a broad amount of researchers and by large imaging studies, such as the HCP and ABCD studies. Nevertheless, even with different voxel sizes our results showed high reliability between the two sequences. Finally, we did not perform any statistical corrections for multiple comparisons in any of our tests. The objective of this paper was to uncover if the two sequences are equivalent, not find the differences. Therefore, using any form of correction for multiple comparisons for our statistical tests would becloud our findings.

## Conclusions

Our results indicate that researchers should adopt or switch to the MPRAGE+PMC sequence in their new studies, especially if there are studying populations with high levels of head motion. Morphometric results obtained from the MPRAGE+PMC sequences are comparable to MPRAGE, especially with the low motion images. Hence, there is no loss if researchers would choose to switch from MPRAGE to MPRAGE+PMC. Additionally, our data from a developmental study, shows that T1’s obtained with PMC have much higher reliability compared to the traditional MPRAGE sequence. However, quality control metrics have shown higher scores for MPRAGE compared to MPRAGE+PMC, mostly caused by increased background noise in the MPRAGE+PMC sequence. Hence, if the population being studied has minimal head motion and the researcher would like to maximize data quality (i.e. SNR), the MPRAGE sequence might be preferred.

## Acknowledgments

We would like to thank the participants and parents for participating in the Healthy Brain Network Initiative. We would also like to thank the many individuals who have provided financial support to the CMI Healthy Brain Network to make the creation and sharing of this resource possible and the partial support the NIH grant R01MH091864 to N.T and M.M.

## Supplementary Material

**Supplementary Table 1 (ST1).** Imaging Session Sequence for the Participant that conducted the regular HBN protocol

| Order | Sequence | Time / Run | Cumulative time |
| --- | --- | --- | --- |
| 1 | localizer_32ch | 0:00:47 | 0:00:47 |
| 2 | ANAT_T1W-RU(MPRAGE) | 0:07:19 | 0:08:06 |
| 3 | cmrr_fMRI_DistortionMap_AP | 0:00:05 | 0:08:11 |
| 4 | cmrr_fMRI_DistortionMap_PA | 0:00:05 | 0:08:16 |
| 5 | cmrr_REST1 | 0:05:08 | 0:13:24 |
| 6 | cmrr_PEER1 | 0:01:56 | 0:15:20 |
| 7 | cmrr_REST2 | 0:05:08 | 0:20:28 |
| 8 | cmrr_MOVIE1 | 0:10:08 | 0:30:36 |
| 9 | cmrr__dMRI_DistortionMap_AP | 0:00:50 | 0:31:26 |
| 10 | cmrr__dMRI_DistortionMap_PA | 0:00:50 | 0:32:16 |
| 11 | cmrr_DKI_018 | 0:07:55 | 0:40:11 |
| 12 | ABCD_T2w_SPC_vNAV_setter | 0:00:00 | 0:40:11 |
| 13 | ABCD_T2w_SPC_vNav | 0:06:35 | 0:46:46 |
| 14 | ABCD_T1w_MPR_vNAV_setter | 0:00:00 | 0:46:46 |
| 15 | ABCD_T1w_MPR_vNAV(MPRAGE+PMC) | 0:07:12 | 0:53:58 |
| 16 | cmrr_PEER3 | 0:01:56 | 0:55:54 |
| 17 | cmrr_MOVIE2 | 0:03:28 | 0:59:22 |

**Supplementary Table 2 (ST2).** Imaging Session Sequence for Participants that conducted the Structural Imaging test-retest protocol. Time in between the end of MPRAGE1 and start of MPRAGE2 sequences is 39 minutes Time in between the end of MPRAGE+PMC1 and start of MPRAGE+PMC2 sequences is 53 minutes The time between the start of MPRAGE+PMC1 and MPRAGE+PMC2 is at least 1:00:30 Time between the start of MPRAGE1 and MPRAGE2 is at least 0:45:49 Time between the start of MPRAGE+PMC1 and MPRAGE2 is at least 0:53:11 Time between HCP1 and MPRAGE+PMC2 0:53:18

| Order | Sequence | Time / Run | Cumulative time |
| --- | --- | --- | --- |
| 1 | localizer_32ch | 0:00:47 | 0:00:47 |
| 2 | ABCD_T1w_MPR_vNAV_setter | 0:00:00 | 0:00:47 |
| 3 | ABCD_T1w_MPR_vNAV(MPRAGE+PMC_1) | 0:07:12 | 0:07:59 |
| 4 | ANAT_T1W-RU(MPRAGE_1) | 0:07:19 | 0:15:18 |
| 5 | cmrr_fMRI_DistortionMap_AP | 0:00:05 | 0:15:23 |
| 6 | cmrr_fMRI_DistortionMap_PA | 0:00:05 | 0:15:28 |
| 7 | cmrr_REST1 | 0:05:08 | 0:20:36 |
| 8 | cmrr_PEER1 | 0:01:56 | 0:22:32 |
| 9 | cmrr_REST2 | 0:05:08 | 0:27:40 |
| 10 | cmrr_MOVIE1 | 0:10:08 | 0:37:48 |
| 11 | cmrr_dMRI_DistortionMap_AP | 0:00:50 | 0:38:38 |
| 12 | cmrr_dMRI_DistortionMap_PA | 0:00:50 | 0:39:28 |
| 13 | cmrr_DKI_018 | 0:07:55 | 0:47:23 |
| 14 | ABCD_T2w_SPC_vNAV_setter | 0:00:00 | 0:47:23 |
| 15 | ABCD_T2w_SPC_vNav | 0:06:35 | 0:53:58 |
| 16 | ANAT_T1W-RU(MPRAGE_2) | 0:07:19 | 1:01:17 |
| 17 | ABCD_T1w_MPR_vNAV_setter | 0:00:00 | 1:01:17 |
| 18 | ABCD_T1w_MPR_vNAV(MPRAGE+PMC_2) | 0:07:12 | 1:08:29 |
| 19 | cmrr_PEER3 | 0:01:56 | 1:10:25 |
| 20 | cmrr_MOVIE2 | 0:03:28 | 1:13:53 |

**Supplementary Table ST3 (ST3).** Partial Correlation scores comparing the difference in cortical thickness measurements (MPRAGE - MPRAGE+PMC) and mean FD across functional scans, controlling for age and sex. Scores that show significant correlation (p<0.05) are color coded, with a positive correlation in red and a negative correlation in blue.

| Area | r-scores | p-values |
| --- | --- | --- |
| ctx_lh_caudalanteriorcingulate | -0.067 | 0.253 |
| ctx_lh_caudalmiddlefrontal | -0.197 | 0.001 |
| ctx_lh_cuneus | -0.028 | 0.627 |
| ctx_lh_entorhinal | -0.151 | 0.010 |
| ctx_lh_fusiform | -0.093 | 0.113 |
| ctx_lh_inferiorparietal | -0.222 | 0.000 |
| ctx_lh_inferiortemporal | -0.087 | 0.139 |
| ctx_lh_insula | -0.131 | 0.025 |
| ctx_lh_isthmuscingulate | -0.039 | 0.501 |
| ctx_lh_lateraloccipital | -0.064 | 0.275 |
| ctx_lh_lateralorbitofrontal | -0.093 | 0.113 |
| ctx_lh_lingual | 0.038 | 0.517 |
| ctx_lh_medialorbitofrontal | -0.067 | 0.250 |
| ctx_lh_middletemporal | -0.062 | 0.293 |
| ctx_lh_paracentral | -0.125 | 0.032 |
| ctx_lh_parahippocampal | 0.009 | 0.874 |
| ctx_lh_parsopercularis | -0.227 | 0.000 |
| ctx_lh_parsorbitalis | -0.222 | 0.000 |
| ctx_lh_parstriangularis | -0.171 | 0.003 |
| ctx_lh_pericalcarine | 0.020 | 0.737 |
| ctx_lh_postcentral | -0.017 | 0.774 |
| ctx_lh_posteriorcingulate | -0.067 | 0.254 |
| ctx_lh_precentral | -0.151 | 0.010 |
| ctx_lh_precuneus | -0.203 | 0.000 |
| ctx_lh_rostralanteriorcingulate | 0.105 | 0.074 |
| ctx_lh_rostralmiddlefrontal | -0.210 | 0.000 |
| ctx_lh_superiorfrontal | -0.305 | 0.000 |
| ctx_lh_superiorparietal | -0.129 | 0.028 |
| ctx_lh_superiortemporal | -0.212 | 0.000 |
| ctx_lh_supramarginal | -0.187 | 0.001 |
| ctx_lh_transversetemporal | -0.168 | 0.004 |
| ctx_rh_caudalanteriorcingulate | -0.011 | 0.848 |
| ctx_rh_caudalmiddlefrontal | -0.173 | 0.003 |
| ctx_rh_cuneus | -0.063 | 0.279 |
| ctx_rh_entorhinal | -0.096 | 0.101 |
| ctx_rh_fusiform | -0.133 | 0.023 |
| ctx_rh_inferiorparietal | -0.130 | 0.026 |
| ctx_rh_inferiortemporal | -0.136 | 0.020 |
| ctx_rh_insula | 0.031 | 0.599 |
| ctx_rh_isthmuscingulate | 0.120 | 0.041 |
| ctx_rh_lateraloccipital | -0.041 | 0.482 |
| ctx_rh_lateralorbitofrontal | -0.090 | 0.126 |
| ctx_rh_lingual | 0.091 | 0.118 |
| ctx_rh_medialorbitofrontal | 0.079 | 0.179 |
| ctx_rh_middletemporal | -0.086 | 0.143 |
| ctx_rh_paracentral | -0.166 | 0.004 |
| ctx_rh parahippocampal | 0.067 | 0.250 |
| ctx_rh_parsopercularis | -0.091 | 0.121 |
| ctx_rh_parsorbitalis | -0.155 | 0.008 |
| ctx_rh_parstriangularis | -0.106 | 0.070 |
| ctx_rh_pericalcarine | 0.064 | 0.279 |
| ctx_rh_postcentral | 0.086 | 0.143 |
| ctx_rh_posteriorcingulate | 0.054 | 0.360 |
| ctx_rh_precentral | -0.123 | 0.035 |
| ctx_rh_precuneus | -0.010 | 0.859 |
| ctx_rh_rostralanteriorcingulate | -0.019 | 0.750 |
| ctx_rh_rostralmiddlefrontal | -0.154 | 0.008 |
| ctx_rh_superiorfrontal | -0.262 | 0.000 |
| ctx_rh_superiorparietal | -0.112 | 0.055 |
| ctx_rh_superiortemporal | -0.107 | 0.067 |
| ctx_rh_supramarginal | -0.098 | 0.095 |
| ctx_rh_transversetemporal | -0.175 | 0.003 |

**Supplementary Table 4 (ST4).** Paired t-test results comparing cortical thickness measurements from MPRAGE and MPRAGE+PMC images, controlling for age, sex, and mean FD during the functional scans. Scores that show significant differences (p<0.05) are color coded, with MPRAGE>MPRAGE+PMC in red and MPRAGE+PMC>MPRAGE in blue.

| Area | t-score | p-value |
| --- | --- | --- |
| ctx_lh_caudalanteriorcingulate | 0.719 | 0.473 |
| ctx_lh_caudalmiddlefrontal | -1.020 | 0.308 |
| ctx_lh_cuneus | 0.934 | 0.351 |
| ctx_lh_entorhinal | -2.182 | 0.030 |
| ctx_lh_fusiform | 0.987 | 0.324 |
| ctx_lh_inferiorparietal | 2.325 | 0.021 |
| ctx_lh_inferiortemporal | -1.626 | 0.105 |
| ctx_lh_insula | 1.348 | 0.179 |
| ctx_lh_isthmuscingulate | 3.169 | 0.002 |
| ctx_lh_lateraloccipital | 1.973 | 0.049 |
| ctx_lh_lateralorbitofrontal | -1.447 | 0.149 |
| ctx_lh_lingual | -0.081 | 0.935 |
| ctx_lh_medialorbitofrontal | 0.943 | 0.347 |
| ctx_lh_middletemporal | -0.614 | 0.540 |
| ctx_lh_paracentral | 1.966 | 0.050 |
| ctx_lh parahippocampal | 1.344 | 0.180 |
| ctx_lh_parsopercularis | 1.426 | 0.155 |
| ctx_lh_parsorbitalis | -0.083 | 0.934 |
| ctx_lh_parstriangularis | 2.571 | 0.011 |
| ctx_lh_pericalcarine | -0.564 | 0.573 |
| ctx_lh_postcentral | -1.736 | 0.084 |
| ctx_lh_posteriorcingulate | 2.273 | 0.024 |
| ctx_lh_precentral | -0.737 | 0.462 |
| ctx_lh_precuneus | 3.955 | 0.000 |
| ctx_lh_rostralanteriorcingulate | 0.417 | 0.677 |
| ctx_lh_rostralmiddlefrontal | 0.461 | 0.645 |
| ctx_lh_superiorfrontal | -0.306 | 0.760 |
| ctx_lh_superiorparietal | 0.387 | 0.699 |
| ctx_lh_superiortemporal | 0.562 | 0.575 |
| ctx_lh_supramarginal | 0.942 | 0.347 |
| ctx_lh_transversetemporal | 1.572 | 0.117 |
| ctx_rh_caudalanteriorcingulate | 1.465 | 0.144 |
| ctx_rh_caudalmiddlefrontal | -0.487 | 0.626 |
| ctx_rh_cuneus | 0.252 | 0.801 |
| ctx_rh_entorhinal | -1.659 | 0.098 |
| ctx_rh_fusiform | -0.790 | 0.430 |
| ctx_rh_inferiorparietal | 0.174 | 0.862 |
| ctx_rh_inferiortemporal | -2.835 | 0.005 |
| ctx_rh_insula | -0.269 | 0.788 |
| ctx_rh_isthmuscingulate | 1.051 | 0.294 |
| ctx_rh_lateraloccipital | 1.677 | 0.095 |
| ctx_rh_lateralorbitofrontal | -0.155 | 0.877 |
| ctx_rh_lingual | -0.875 | 0.382 |
| ctx_rh_medialorbitofrontal | 1.542 | 0.124 |
| ctx_rh_middletemporal | -0.803 | 0.423 |
| ctx_rh_paracentral | 2.497 | 0.013 |
| ctx_rh parahippocampal | -0.166 | 0.869 |
| ctx_rh_parsopercularis | 0.325 | 0.746 |
| ctx_rh_parsorbitalis | -0.076 | 0.939 |
| ctx_rh_parstriangularis | 1.059 | 0.291 |
| ctx_rh_pericalcarine | -0.129 | 0.898 |
| ctx_rh_postcentral | -1.929 | 0.055 |
| ctx_rh_posteriorcingulate | 2.305 | 0.022 |
| ctx_rh_precentral | -1.790 | 0.075 |
| ctx_rh_precuneus | 2.060 | 0.040 |
| ctx_rh_rostralanteriorcingulate | 2.589 | 0.010 |
| ctx_rh_rostralmiddlefrontal | 0.521 | 0.603 |
| ctx_rh_superiorfrontal | 0.376 | 0.707 |
| ctx_rh_superiorparietal | -0.359 | 0.720 |
| ctx_rh_superiortemporal | 0.210 | 0.834 |
| ctx_rh_supramarginal | -0.685 | 0.494 |
| ctx_rh_transversetemporal | 2.924 | 0.004 |

## Supplementary Figures

**Supplementary Figure 1 (SF1).**
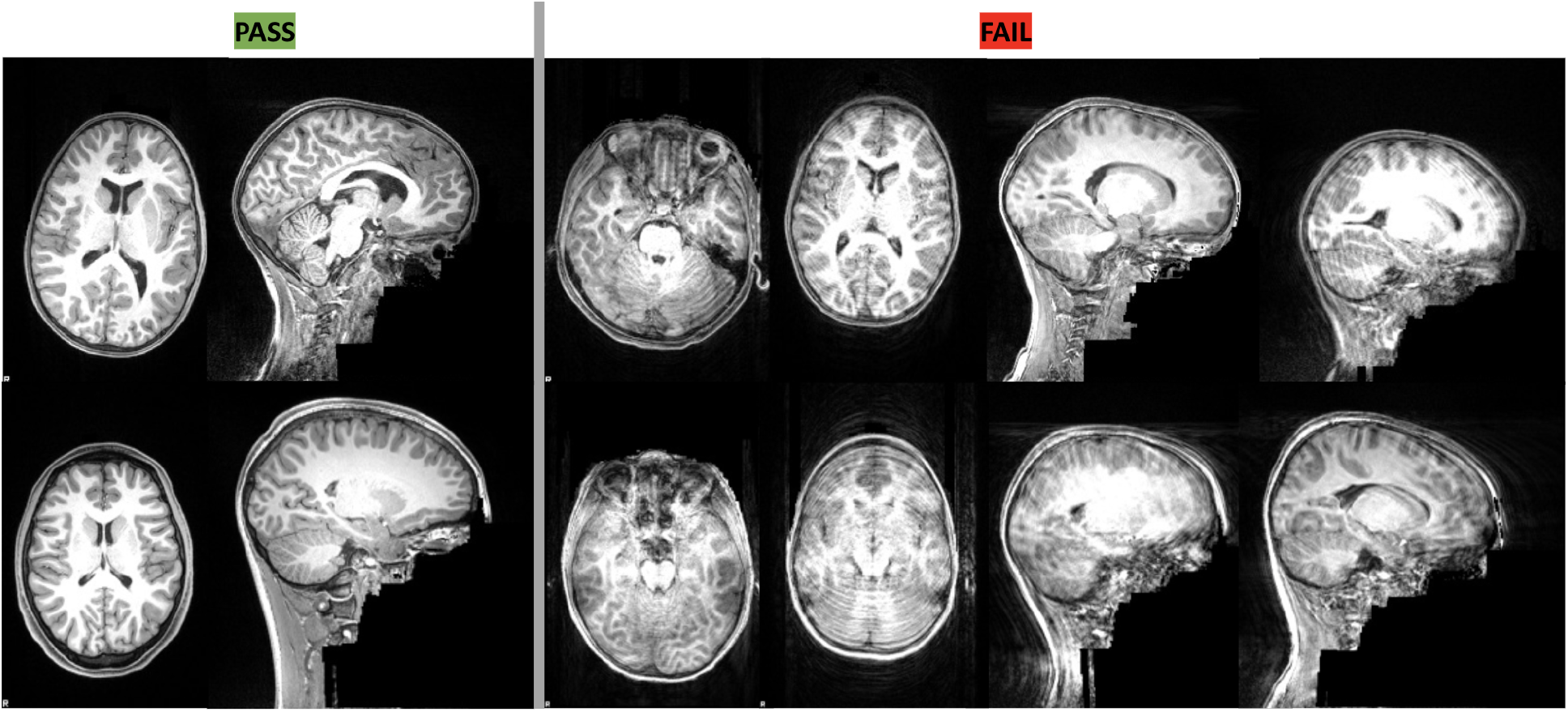
Example figures used for training raters for visual quality control.

**Supplementary Figure 2 (SF2).**
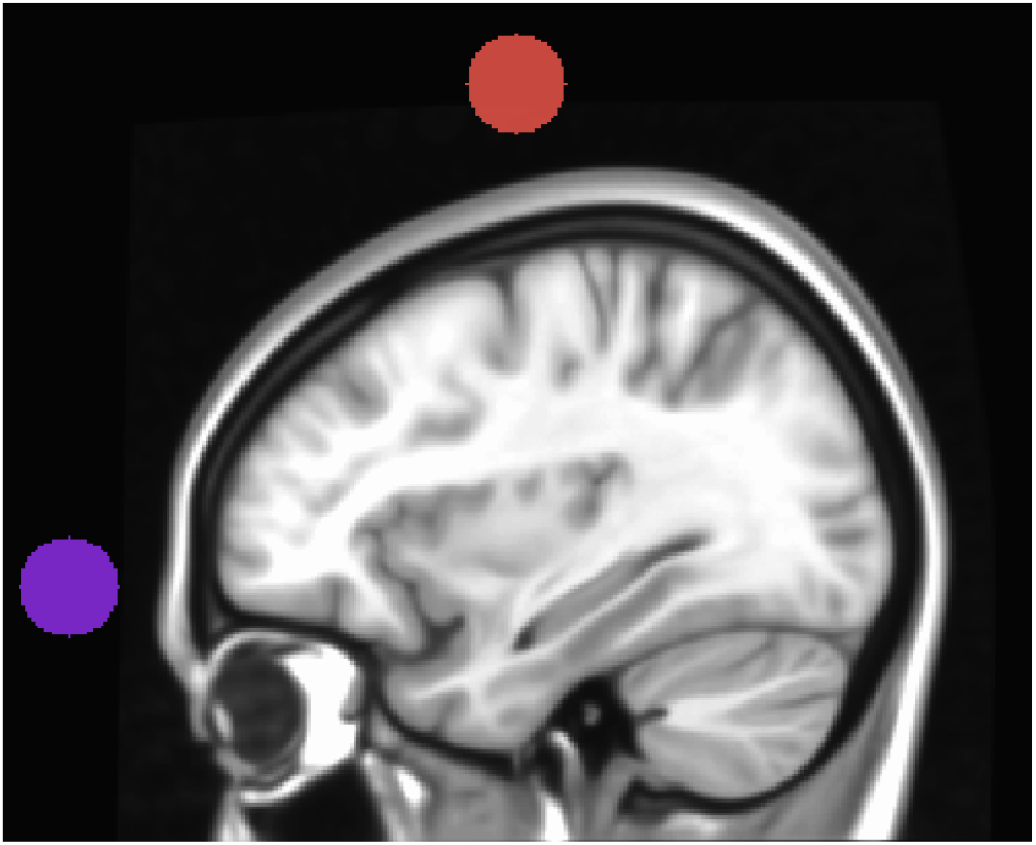
Location of circles for calculating Anterior-to-Superior Ratio (ASR).

**Supplementary Figure 3(SF3).**
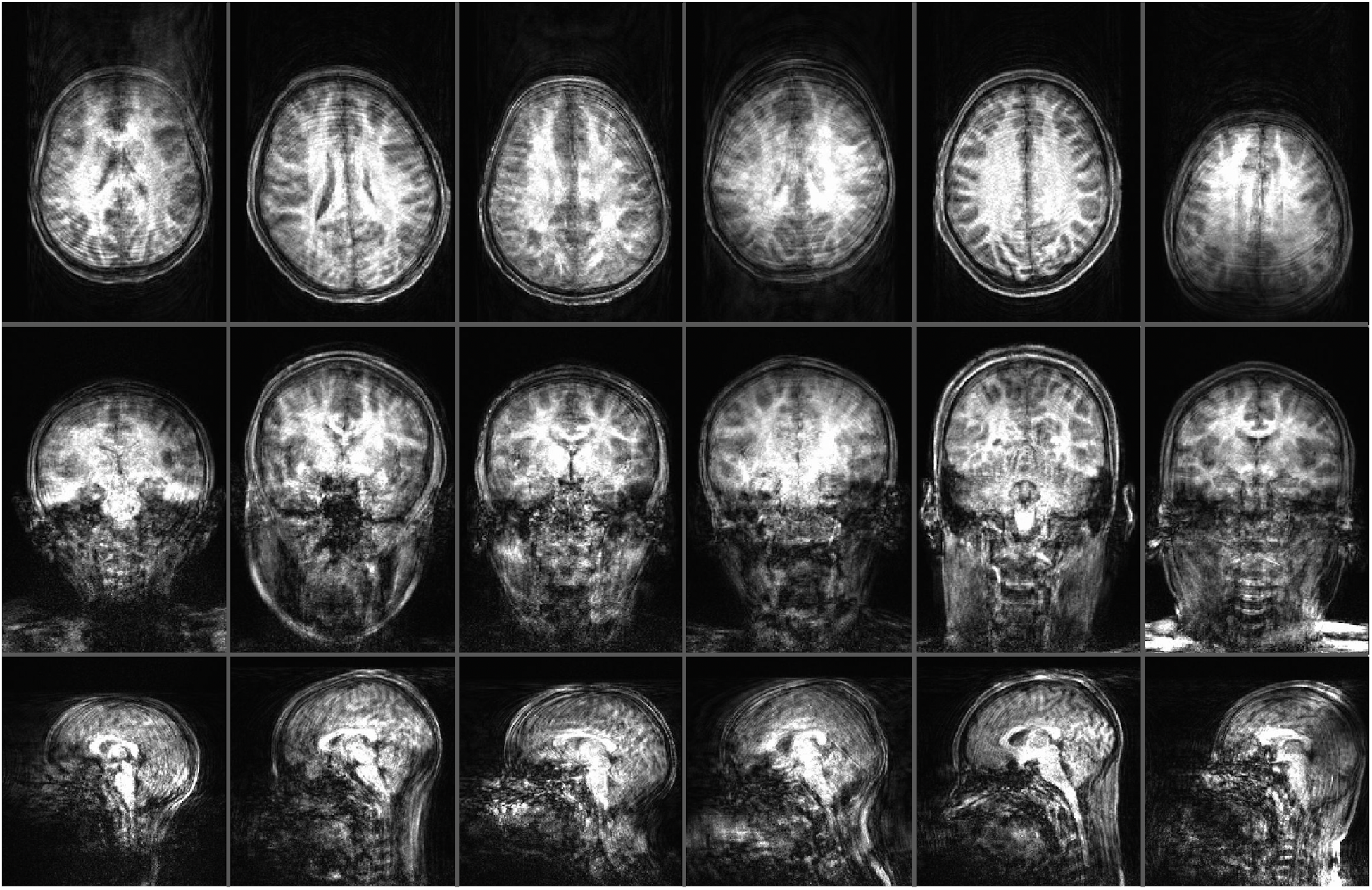
Images with high motion for MPRAGE+PMC runs

**Supplementary Figure (SF4).**
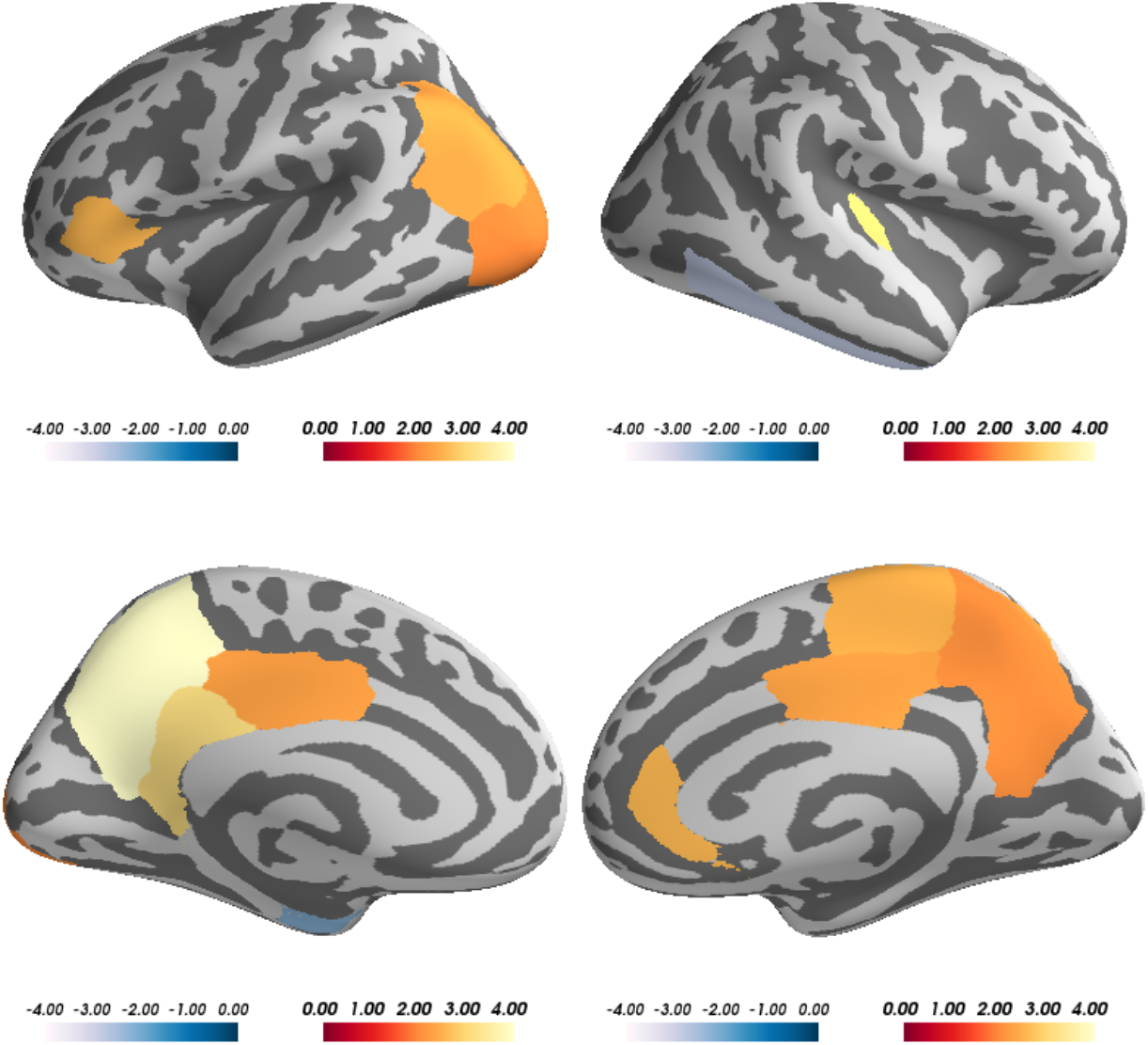
Paired t-test results comparing cortical thickness measurements from MPRAGE and MPRAGE+PMC images. Regions of the brain with significant difference (p<0.05) are color-coded based on t-scores from the paired t-test, where warm colors represent MPRAGE > MPRAGE+PMC and cold colors represent MPRAGE+PMC > MPRAGE.

**Supplementary Figure 5 (SF5).**
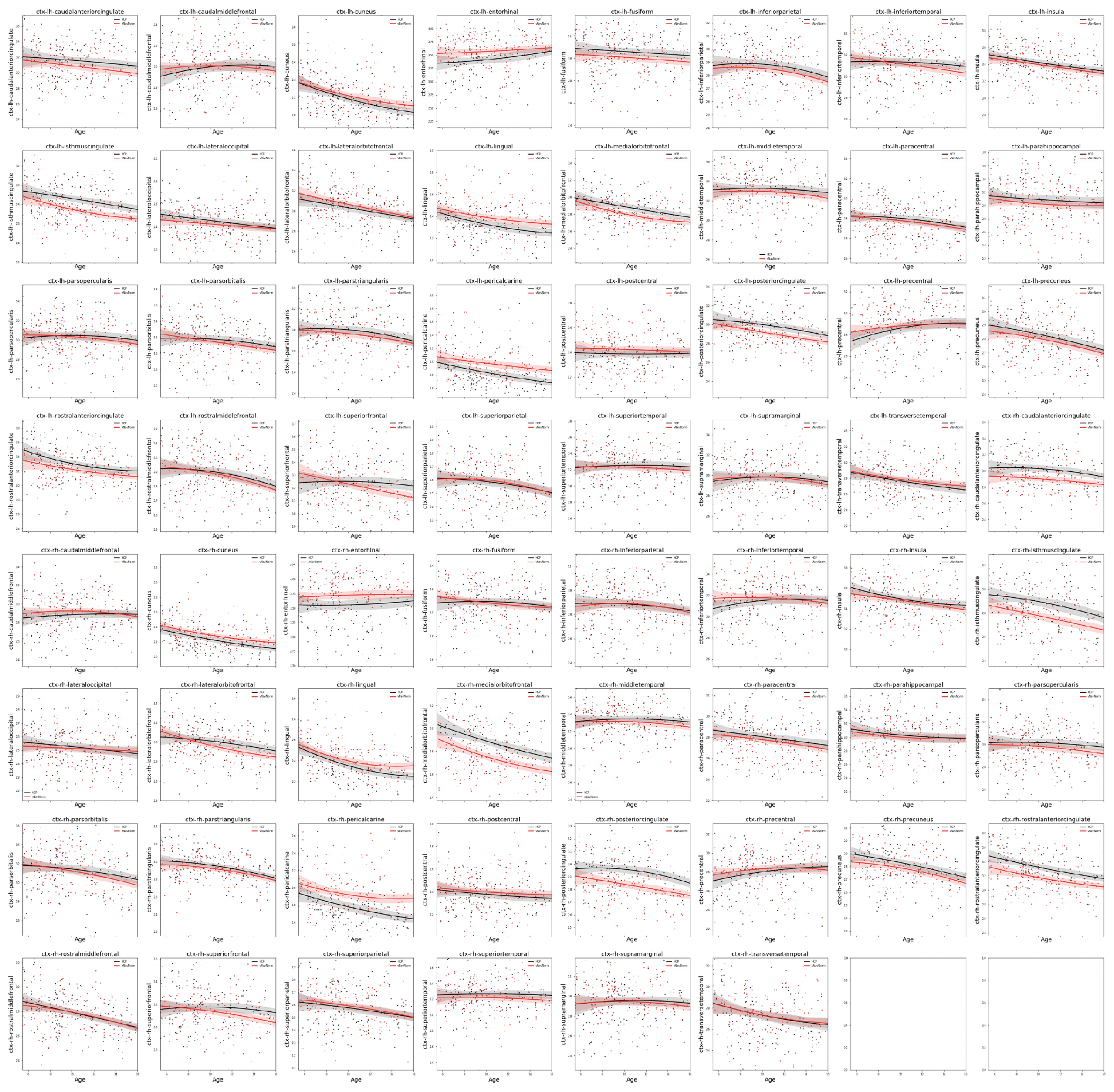
Cortical Thickness Development curves for Males.

**Supplementary Figure 6 (SF6).**
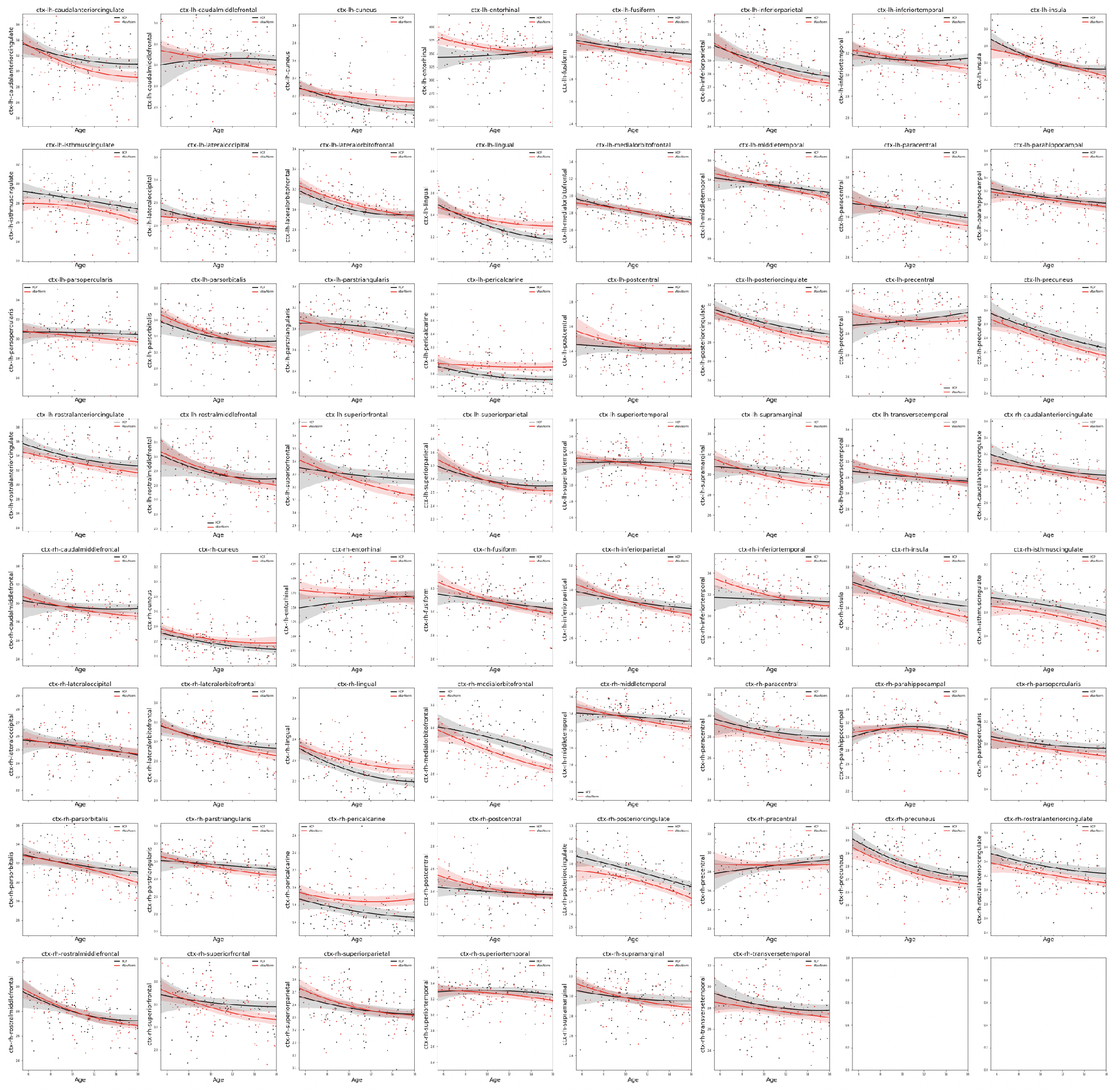
Cortical Thickness Development curves for Females.

**Supplementary Figure 7 (SF7).**
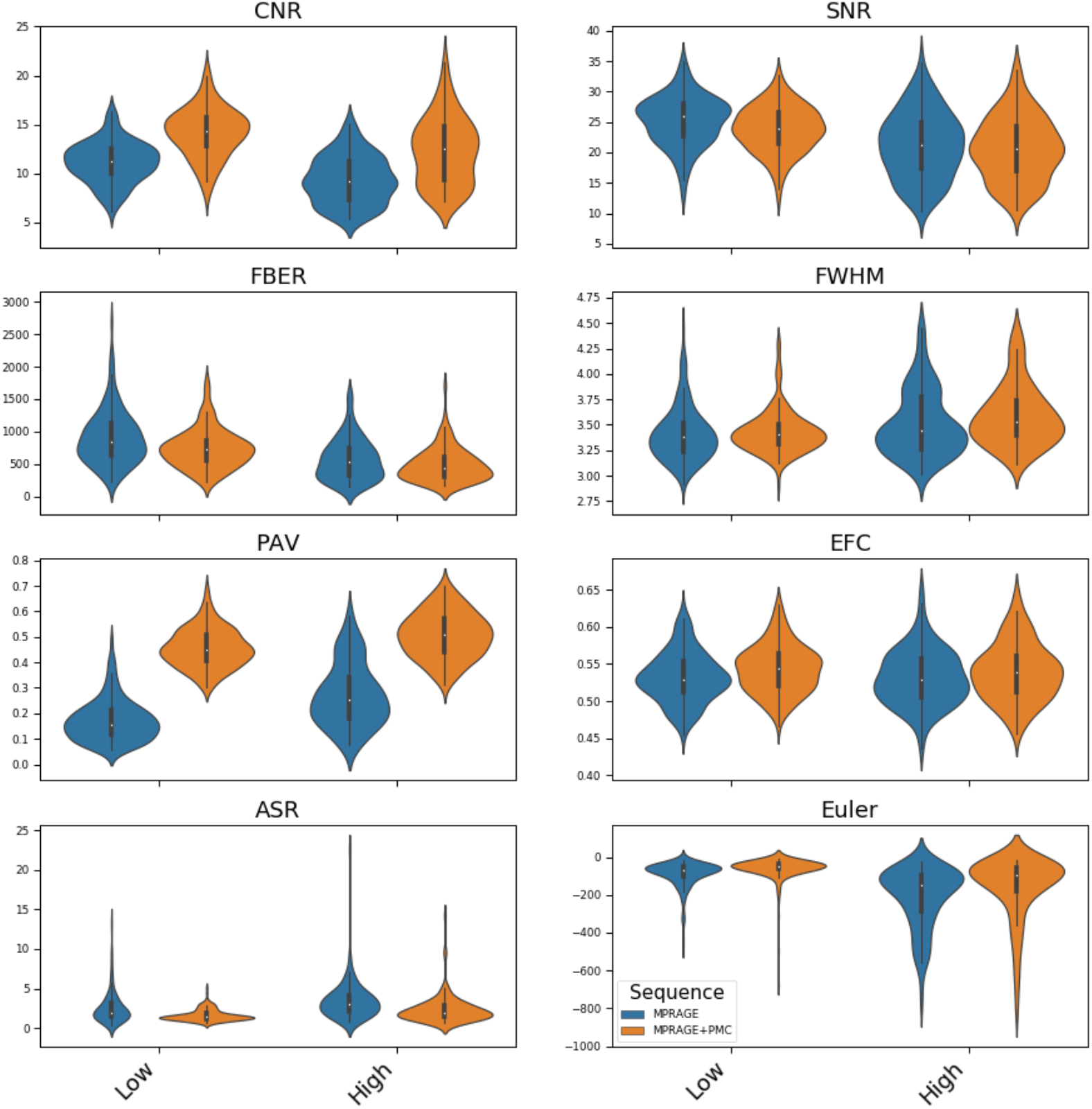
Quality control metrics for the MPRAGE and MPRAGE+PMC images across 287 participants, separated by low and high movers.

**Supplementary Figure 8 (SF8).**
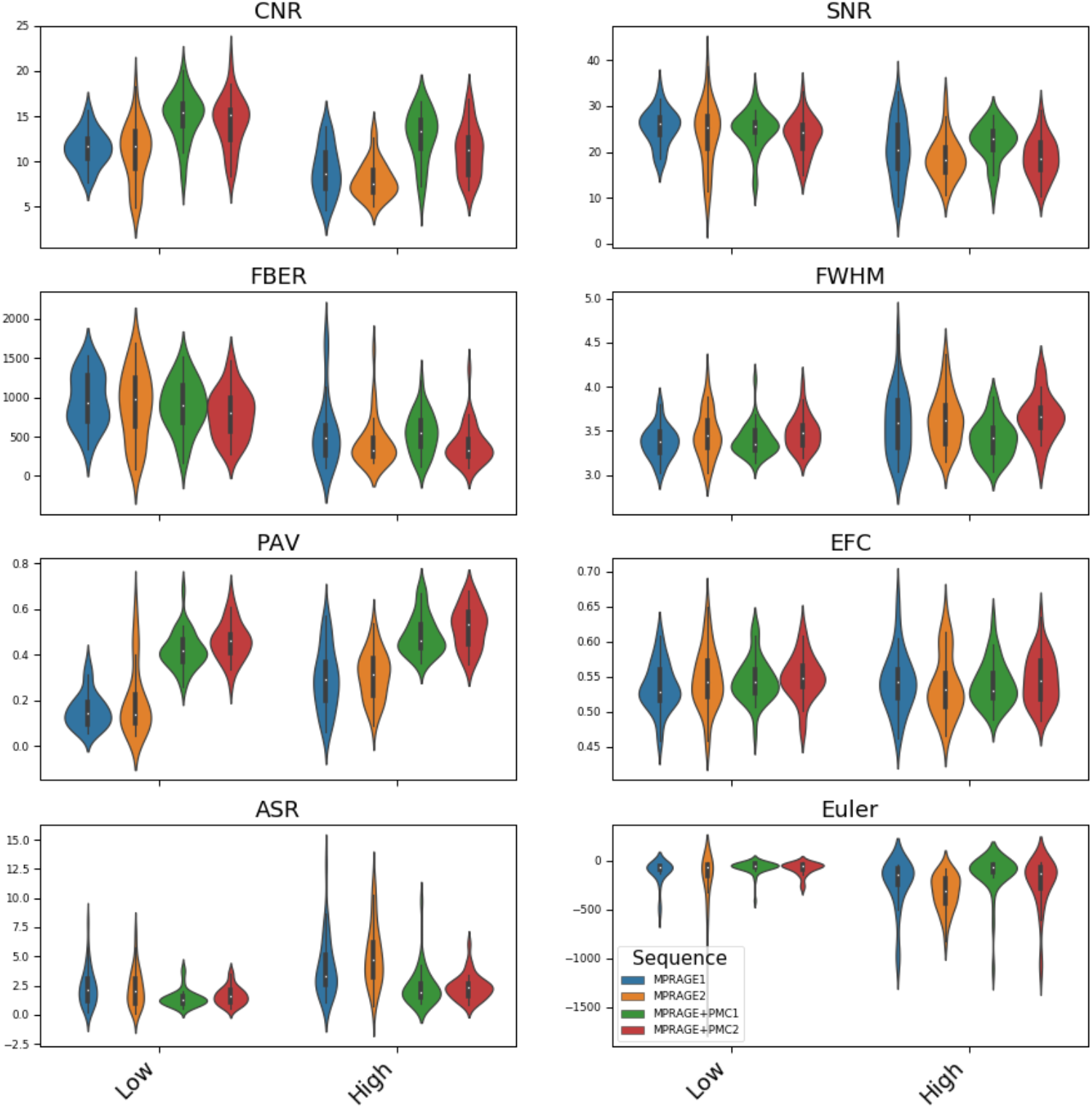
Quality Control Metrics for the test-retest group.

## Footnotes

1 MPRAGE is the sequence used by Siemens MRIs and the equivalent of this sequence for GE machines is the 3-D Fast SPGR and for Philips is its 3D TFE

2 MPRAGE is the recommended sequence to be used by Freesurfer

3 It is possible to directly obtain the RMS motion estimation scores from the Dicom files using a python function available here: https://github.com/MRIMotionCorrection/parse_vNav_Motion. However, we choose to recalculate the motion estimation parameters with QAP to maintain consistency with the functional data.

4 Peer (Peer Eye Estimation Regression) is a short (<2 minutes) functional run to calibrate an fMRI-based eye-tracking algorithm. See (Son et al. 2019) for more details. In the initial HBN protocol, there was a peer2 run which was later dropped given time constraints and no need to have 3 calibration runs.

